# Enhancing antibody responses by multivalent antigen display on thymus-independent DNA origami scaffolds

**DOI:** 10.1101/2022.08.16.504128

**Authors:** Eike-Christian Wamhoff, Larance Ronsard, Jared Feldman, Grant A. Knappe, Blake M. Hauser, Anna Romanov, Evan Lam, Kerri St. Denis, Julie Boucau, Amy K Barczak, Alejandro B. Balazs, Aaron Schmidt, Daniel Lingwood, Mark Bathe

## Abstract

Multivalent antigen display is a well-established principle to enhance humoral immunity. Protein-based virus-like particles (VLPs) are commonly used to spatially organize antigens. However, protein-based VLPs are limited in their ability to control valency on fixed scaffold geometries and are thymus-dependent antigens that elicit neutralizing B cell memory themselves, which can distract immune responses. Here, we investigated DNA origami as an alternative material for multivalent antigen display in vivo, applied to the receptor binding domain (RBD) of SARS-CoV-2 that is the primary antigenic target of neutralizing antibody responses. Icosahedral DNA-VLPs elicited neutralizing antibodies to SARS-CoV-2 in a valency-dependent manner following sequential immunization in mice, quantified by pseudo-and live-virus neutralization assays. Further, induction of B cell memory against the RBD required T cell help, but the immune sera did not contain boosted, class-switched antibodies against the DNA scaffold. This contrasted with protein-based VLP display of the RBD that elicited B cell memory against both the target antigen and the scaffold. Thus, DNA-based VLPs enhance target antigen immunogenicity without generating off-target, scaffold-directed immune memory, thereby offering a potentially important alternative material for particulate vaccine design.

## Introduction

Multivalent display of antigens on virus-like particles (VLPs) can markedly improve the immunogenicity of subunit vaccines^1-3^. Nanoparticulate vaccines with diameters between 20 and 200 nm ensure efficient trafficking to secondary lymphoid organs, and particle diameters below 50 nm mitigate undesired retention at the injection site and promote the penetration of B cell follicles^4,5^. In secondary lymphoid organs, multivalency promotes B cell receptor (BCR) crosslinking and signaling as well as BCR-mediated antigen uptake, thereby driving B cell activation and humoral immunity^6-13^.

The importance of BCR signaling for the generation of antibody responses was initially recognized for thymus-independent (TI) antigens, particularly of the TI-2 class^14-16^. The multivalent display of these non-protein antigens induces BCR crosslinking in the absence of T cell help. The resultant antibody responses proceed through extrafollicular B cell pathways, with limited germinal center (GC) reactions, affinity maturation, and induction of B cell memory^17,18^. Multivalent antigen display also enhances BCR-mediated responses to thymus-dependent (TD) antigens, namely proteins^8,9^. In this context, follicular T cell help enables GC reactions to generate affinity-matured B cell memory that can be boosted or recalled upon antigen reexposure^19-21^. Consequently, the nanoscale organization of antigens represents a well-established vaccine design principle not only for TI antigens, but also to elicit humoral immunity through the TD pathway^1-3^.

Leveraging this design principle, protein-based virus-like particles (P-VLPs) have emerged as an important material platform for multivalent subunit vaccines^22-38^. P-VLPs enable the rigid display of TD antigens and have been used to investigate the impact of valency on B cell activation in vivo, suggesting early B cell activation and downstream humoral immune responses are improved substantially for some antigens as valency increases^8-10^. However, control over antigen valency in P-VLPs is constrained to the constituent self-assembled protein scaffold subunits, rendering the investigation of antigen valency on humoral immunity challenging without simultaneously altering scaffold size, geometry, and protein composition^9,10^. Alternatively, if a constant protein scaffold geometry is used, then current approaches are limited to stochastically-controlled antigen valency and spatial positioning^8,29,30,38^. Further, protein-based scaffolds are TD antigens that elicit humoral immunity themselves^38-40^. This potentially distracts antibody responses from the target antigens of interest^41,42^, and might also lead to immune imprinting^43^ or original antigenic sin (OAS)^44,45^ in which off-target, immunodominant epitopes distract from target epitopes of interest in generating B cell memory. Finally, scaffold-directed immunological memory may also result in antibody-dependent clearance of the vaccine material, thereby limiting sequential or diversified immunizations with a given P-VLP ^46,47^.

To overcome these limitations of protein-based materials, here we sought to design rigid, TI scaffolds of fixed geometry and size to display target TD antigens of varying valency. Fixing scaffold geometry would isolate the impact of valency on promoting antibody responses, whereas use of a TI scaffold would promote focusing of the antibody response on the target, TD antigen of interest while confining scaffold-directed B cell responses to the non-boostable, TI pathway that is devoid of immunological memory^48,49^. This approach may therefore avoid the off-target, distracting antibody responses that affect P-VLPs^40,50^ while retaining the ability to enhance humoral immunity through multivalent antigen display. Wireframe DNA origami provides unique access to such rationally designed VLPs of the optimal 50 nm size-scale, with scaffold-independent control over valency of antigen display^51-56^. While we and others have leveraged this DNA-based material platform in vitro to probe the nanoscale parameters of IgM recognition^57^ and BCR signaling in reporter B cell lines^58^, the functional properties of this material remained to be investigated in vivo where complex processes including particle trafficking, T cell help, and scaffold degradation mediated by endonucleases are present.

To investigate the ability of these materials to elicit functional immune responses in vivo, DNA-VLPs functionalized with the SARS-CoV-2 receptor binding domain (RBD) derived from the spike glycoprotein, a key target for eliciting neutralizing antibodies against the virus^59-62^, were fabricated with varying valency, validated in vitro, and subsequently evaluated in vivo. This nanoparticulate vaccine displayed enhanced binding to ACE2-expressing cells in vitro, and induced BCR signaling in a valency-dependent manner in vitro consistent with previous findings^58,63,64^. Following sequential vaccine immunizations in mice, we observed valency-dependent enhancement of RBD-specific IgG responses and B cell memory recall in vivo using this DNA-based scaffold material. Functionally, neutralization of the wildtype, Wuhan strain of SARS-CoV-2 was found to be more efficient for antibodies elicited by DNA-VLPs compared with those elicited by monomeric RBD alone. However, immunized sera did not contain boosted, DNA-specific antibodies, and antibody enhancement was not observed in *Tcra*^−/−^ mice, demonstrating that boosting of RBD-specific antibodies went through the TD pathway even though antigens were presented on TI scaffolds. This contrasted with a commonly used protein-based vaccine scaffold that generated and boosted specific B cell memory to both the target, RBD antigen, and the scaffold protein material. Taken together, our findings suggest that DNA origami-based materials can be used to enhance humoral immunity in vivo using multivalent antigen display via focusing of antibody responses on target antigens without the generation of distracting B cell memory against the VLP scaffold itself.

## Results and Discussion

SARS-CoV-2 displays trimeric spike glycoproteins on a ∼100 nm diameter scaffold^65^, where each glycoprotein monomer contains the receptor-binding domain (RBD) for engaging the ACE2 receptor mediating viral uptake, thereby rendering RBD a key target for neutralizing antibody responses^59-62^. To ensure optimal trafficking of our vaccine platform, which requires particle diameters smaller than 50 nm^5^, we computationally designed and fabricated an icosahedral DNA-VLP with 30 conjugation sites and approximately 34 nm scaffold diameter to display the RBD^52,54^. A covalent *in situ* functionalization strategy employing strain-promoted azide-alkyne cycloaddition (SPAAC) chemistry was used to ensure rigid, irreversible antigen attachment to the DNA scaffold (**Figure 1A**)^53^, unlike previous work that used a reversible, hybridization strategy^58^. Towards this end, we synthesized 30 oligonucleotide staples bearing triethylene glycol (TEG)-DBCO groups at their 5’ ends to assemble DNA-VLPs symmetrically displaying 1x, 6x or 30x DBCO groups on their exterior (**Figure S1, Table S1 to S3**). Employing a reoxidation strategy, the monomeric RBD was selectively modified at an engineered C-terminal Cys with an SMCC-TEG-azide linker to yield RBD-Az which was subsequently incubated with DBCO-bearing DNA origami to fabricate **I52-1x, -6x**, and **-30x** (**Figures 1B and S2, Note S1**). The optimization of reaction conditions yielded near-quantitative functionalization efficiency of conjugation sites as determined by denaturing, reversed-phase HPLC^53^ and Trp fluorescence^58^ (**Figures 1C and S3**). The monodispersity of purified DNA-VLPs was validated by dynamic light scattering (DLS) (**Figure 1D**). Analysis of **I52-30x** via negative-strain transmission electron microscopy (TEM) validated structural integrity of the DNA origami with the desired total size of approximately 45 nm (**Figure 1E and S4**). While the icosahedral geometry could not be fully resolved, presumably due to accumulation of uranyl formate in the interior of the DNA origami, antigens were clearly visible and organized symmetrically.

**Figure 1.**
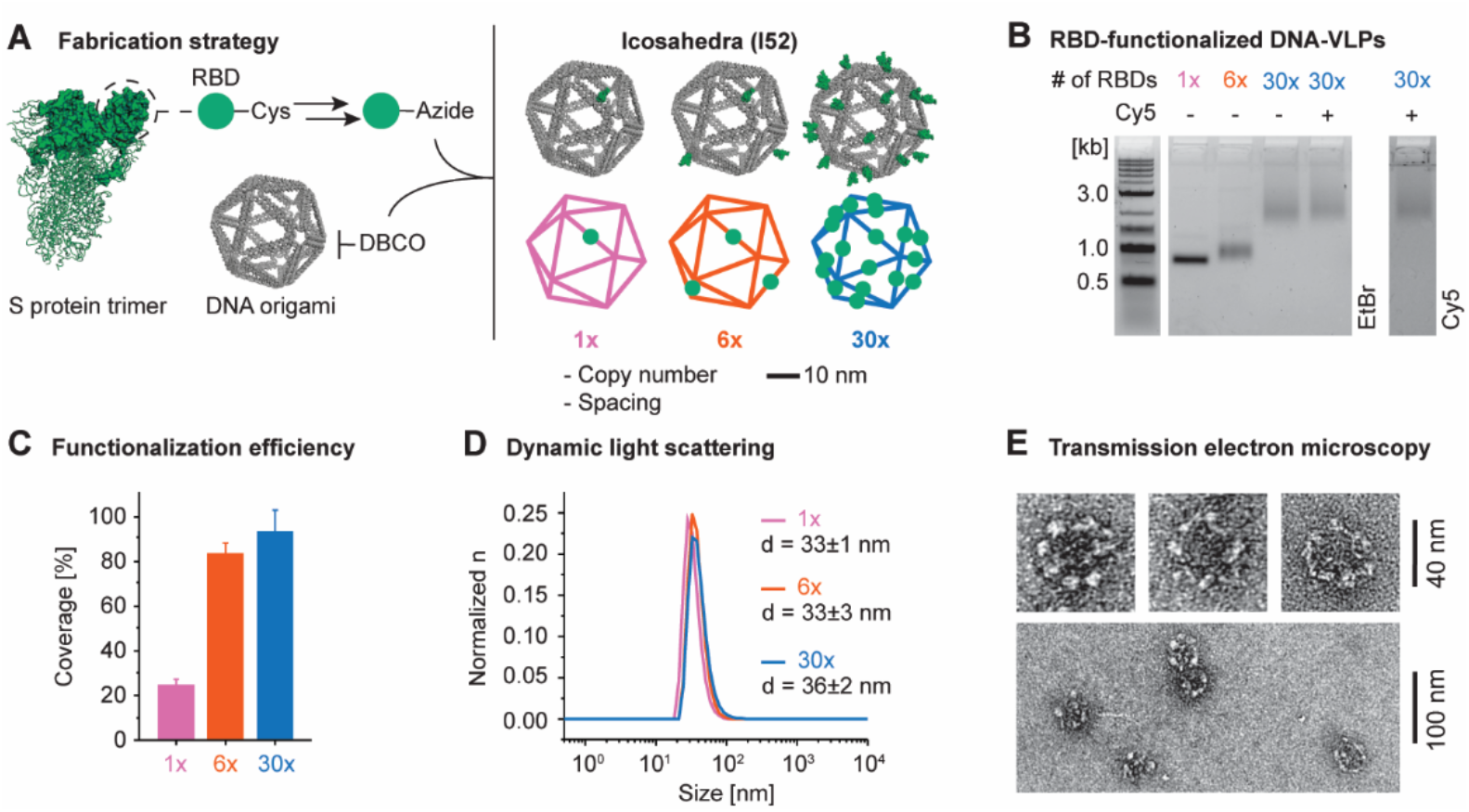
Design and synthesis of DNA-VLPs covalently displaying the SARS-CoV-2 RBD. **(A)** Recombinant RBD bearing an additional Cys residue at the C-terminus was expressed. The C-terminal Cys was selectively labeled with and SMCC-TEG-azide linker and subsequently conjugated to DBCO-bearing DNA-VLPs. The icosahedral DNA origami objects of approximately 50 nm diameter displaying 1, 6 and 30 copies of the RBD were fabricated. **(B)** Agarose gel electrophoresis (AGE) shows the gel shift due to increasing RBD copy number as well as low polydispersity of the VLPs samples after purification. An additional VLP bearing 5 copies of Cy5 was produced for ACE2-binding flow cytometry experiments. **(C)** The coverage of the DNA-VLPs with RBD was quantified via Trp fluorescence. **(D)** Dynamic light scattering (DLS) was used to assess the dispersity of functionalized VLP samples. Representative histograms are shown. **(E)** Transmission electron micrographs (TEM) of **I52-30x** were obtained by negative staining using 2% uranyl formate and validate the symmetric nanoscale organization of antigens. Coverage values were determined from n = 3 biological replicates for **I52-1x** and from n = 6 biological replicates for **I52-6x** and **I52-30x**. Diameters were determined from 3 technical replicates. Error bars represent the standard error of the mean.

To investigate the binding activity of RBD-Az before and after conjugation to DNA-VLPs, we conducted flow cytometry experiments with ACE2-expressing HEK293 cells (**Figure 2A**). Initially, monovalent binding of wild-type RBD and fluorophore-labeled RBD-Cy5, obtained by selectively labeling the azide, was compared (**Figure 2B and C**). The RBD constructs were incubated at 200 nM with the HEK293 cells and bound antigen was detected using the previously described anti-RBD antibody CR3022^62^. These experiments revealed comparable binding between the two constructs, demonstrating preservation of binding activity of the receptor binding motif (RBM) in RBD and the viability of the reoxidation strategy for selective labeling of the terminal Cys (**Figure S2, Note S1**).

**Figure 2.**
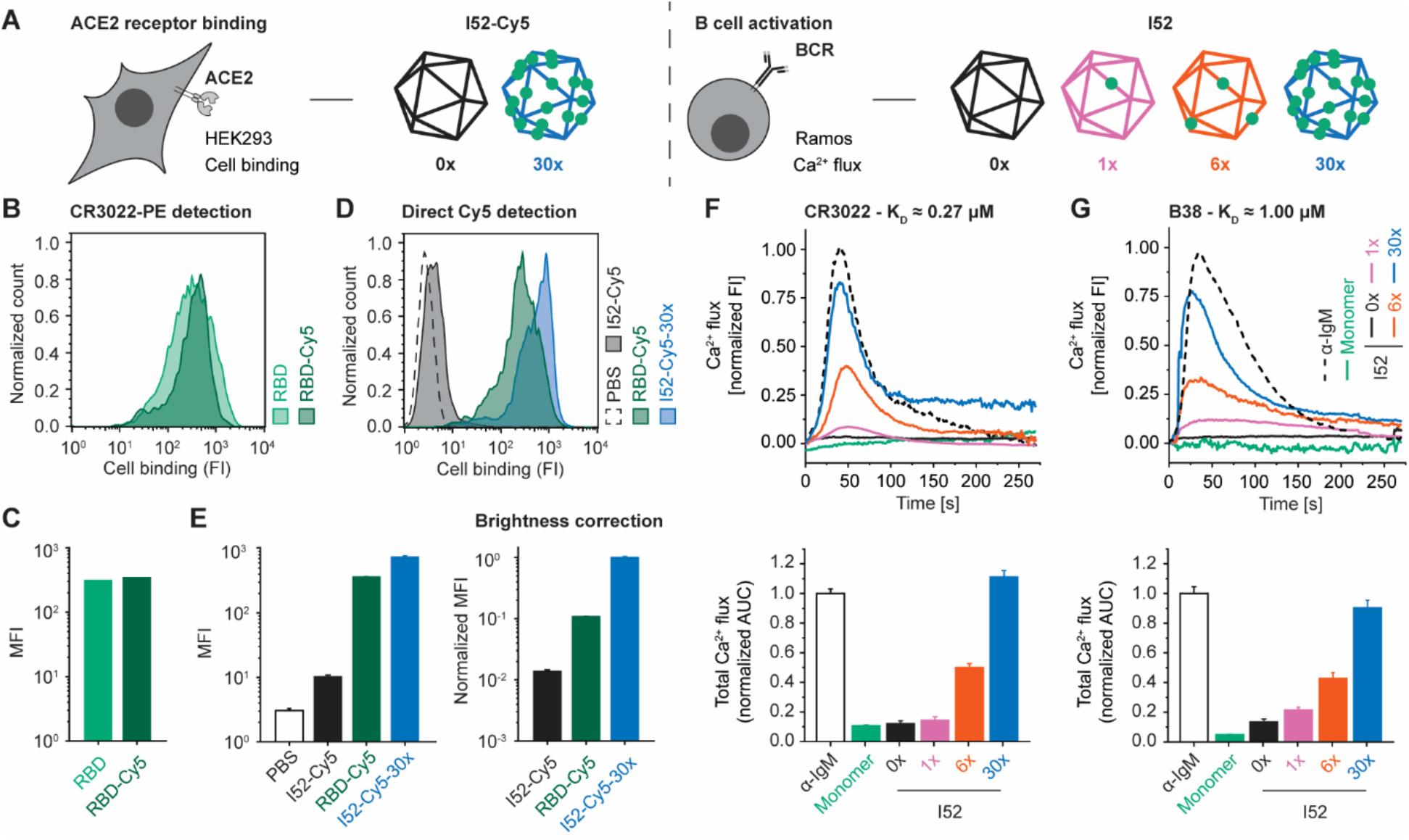
In vitro activity of RBD-functionalized DNA-VLPs. **(A)** An overview of the in vitro activity assays and corresponding DNA-VLPs is shown. **(B and C)** ACE2-expressing HEK293 cells were incubated with 200 nM RBD. Binding was detected in flow cytometry experiments using PE-labeled CR3022 and a PE-labeled secondary antibody, demonstrating preserved binding activity for chemically modified RBD-Cy5 compared to wild-type RBD. **(D and E)** Incubation with Cy5-labeled **I52-Cy5-30x** at 100 nM RBD revealed enhanced binding compared to RBD-Cy5 due to multivalency effects. No unspecific binding for non-functionalized **I52-Cy5** was observed. The brightness of Cy5-labeled **I52-Cy5-30x** (5 Cy5 per 30 RBDs) and RBD-Cy5 (1 Cy5 per 1 RBD) was quantified experimentally (**Figure S4**) and MFI values were corrected accordingly. **(F and G)** Ramos B cells expressing the BCRs C3022 and B38 were incubated with α-IgM, wild-type RBD or RBD-functionalized DNA-VLPs at 30 nM RBD. Ca^2+^ flux in response to RBD incubation was assayed using Fura Red. Representative fluorescence intensity curves are shown (top). Total Ca^2+^ flux was quantified via the normalized AUC, revealing robust activation of BCR-expressing Ramos B cells by functionalized DNA-VLPs (bottom). No stimulation was observed for wild-type RBD or for non-functionalized **I52-0x**. Representative histograms are shown for ACE2 binding assays and MFI values were determined from n = 3 biological replicates. Normalized AUC values were determined from n = 3 biological replicates. Error bars represent the standard error of the mean.

Next, we explored whether multivalent RBD display using DNA-VLPs would result in increased avidity. Two additional fluorophore-labeled DNA-VLPs, **I52-Cy5-30x** and **I52-Cy5**, were fabricated to allow for direct detection of binding (**Figure 1B and S1**). Binding of **I52-Cy5-30x** was significantly enhanced compared to monomeric RBD-Cy5, while no binding was observed for the **I52-Cy5** (**Figure 2D and 2E**). When correcting for Cy5 brightness per RBD, **I52-Cy5-30x** displayed approximately one order of magnitude higher median fluorescence intensity compared with monomeric RBD-Cy5, likely due to avidity effects in multivalent DNA-VLP binding to the cognate ACE2 receptor.

We then evaluated the impact of RBD-functionalized DNA-VLPs on BCR signaling using a previously described Ca^2+^ flux assay (**Figure 2A**)^63^. Specifically, Ramos B cell lines expressing the somatic CR3022 or B38 antibodies were established^62,66^ and BCR signaling was validated by incubation with anti-IgM antibody. At 30 nM antigen concentration, monomeric RBD did not elicit B cell activation in vitro (**Figure 2F and 2G**). In contrast, incubation of the Ramos B cells with multivalent DNA-VLPs at the same antigen concentration resulted in efficient BCR signaling. We further observed valency-dependent increases in total Ca^2+^ flux for both cell lines with **I52-30x** more potent than **I52-6x**. CR3022 (K_D_ = 0.27 μM, **Figure 2F**) and B38 (K_D_ = 1.00 μM, **Figure 2G**) bind distinct RBD epitopes with moderate monovalent affinity as reported for the corresponding Fab fragments^67^. Despite this four-fold difference in affinity, we observed comparable total BCR signaling relative to the IgM control for all functionalized DNA-VLPs, consistent with previously described avidity effects at the B cell surface^68^. We thereby concluded that our DNA-VLPs efficiently bound and induced signaling by RBD-specific BCRs in a valency-dependent manner, analogous to previous studies using similar assays to evaluate protein- and DNA-scaffolded multivalent subunit vaccines^58,63,64,69-74^.

To investigate whether RBD-functionalized DNA-VLPs activated B cells in vivo to induce antibody responses, C57BL/6 mice were sequentially immunized intraperitoneally with monomeric RBD, **I52-6x**, or **I52-30x** at doses equivalent to 7.5 μg RBD with Sigma adjuvant (**Figure 3A and S3**). Serum IgG responses against the RBD were monitored throughout this regimen using ELISA and correlated with in vitro BCR signaling findings (**Figure 3B and S5**). Following boost 1, we observed an approximately 130-fold increase in endpoint dilutions for **I52-30x** over monomeric RBD. In contrast, **I52-6x** elicited comparable antibody responses with respect to monomeric RBD following each boost, suggesting that a higher minimum antigen copy number is needed to enhance B cell responses in vivo compared with in vitro (**Figure 2F and 2G**), which may be due to differences in trafficking or degradation rates, for example, between **I52-6x** and **I52-30x**. Overall, IgG titers increased for monomeric RBD, **I52-6x**, and **I52-30x** following boost 2, converging to similar endpoint dilutions. These findings of earlier and stronger boosting of IgG titers and efficient B cell memory recall elicited by the **I52-30x** are hallmarks of multivalent versus monomeric subunit vaccines^23,24^, consistent with enhanced IgG titers elicited by P-VLPs of increasing valency^8-10^.

**Figure 3.**
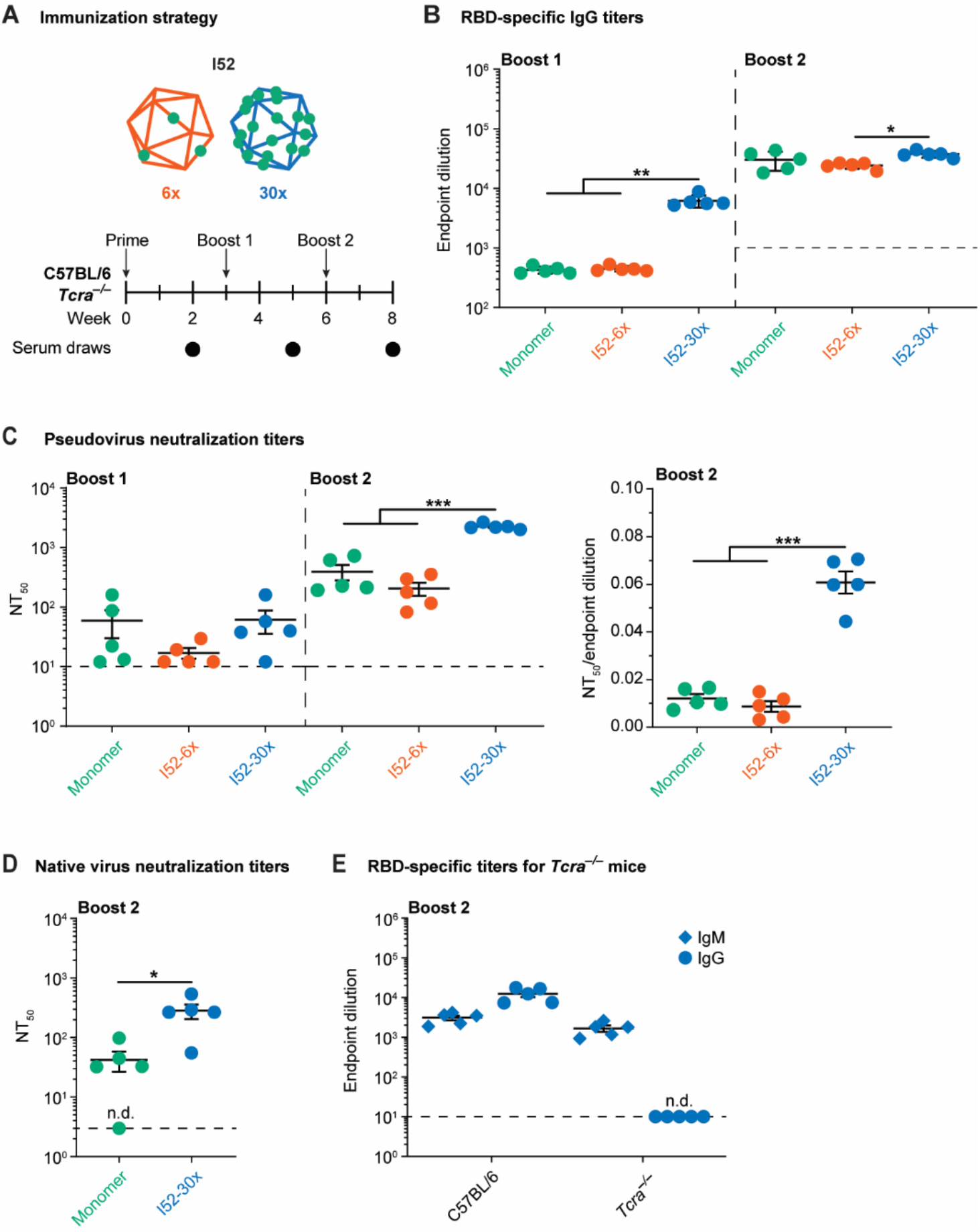
RBD-specific antibody responses to functionalized DNA-VLPs. **(A)** Mice were sequentially immunized with monomeric RBD and RBD-functionalized DNA-VLPs of varying copy number. **(B)** RBD-specific IgG endpoint dilutions revealed enhanced antibody responses for **I52-30x** compared to both monomeric RBD and **I52-6x. (C)** Serum neutralization titers expressed as NT_50_ values against pseudoviruses modeling the wild-type, Wuhan strain. **(D)** Neutralization of native Wuhan SARS-CoV-2. **(E)** IgM and IgG titers of RBD-specific antibodies elicited in *Tcra*^−/−^ and wild-type mice after sequential immunization with **I52-30x**. N=5 animals were used in each experimental group. One-way ANOVA was performed followed by Dunnett’s T3 multiple comparison test at α = 0.05 for **(B)** and **(C)**. Welch’s t-test was performed at α = 0.05 for **(D)**. Error bars represent the standard error of the mean. Non-responder mice (denoted as n.d. = not detectable) were not considered for statistical analysis.

To quantify the quality of antibody responses generated by DNA-VLPs, valency-dependent enhancement of RBD-specific antibody responses was interrogated using viral neutralization assays (**Figure 3C, 3D and S6**). In both pseudo- and native virus neutralization assays, efficient neutralization of the wild-type, Wuhan strain of SARS-CoV-2 resulted from DNA-VLPs. However, across both the pseudo- and native virus assays, **I52-30x** elicited IgG titers with significantly higher neutralization potency compared with both monomeric RBD and **I52-6x**, indicating that the higher valency, 30x-RBD DNA-VLP produced significantly superior quality antibodies, which have been correlated with improved patient outcomes^75-77^.

To confirm that the boosted, RBD-specific IgG titers were achieved via the TD route, we compared the same sequential immunization regime in wild-type C57BL/6 versus *Tcra*^−/−^ mice that lack functional TCRs^78^ (**Figure 3A**). **I52-30x** elicited robust IgM responses in both genotypes, whereas RBD-specific IgG titers were not boosted in *Tcra*^−/−^ mice (**Figure 3E**). Hence, IgG boosting following sequential immunization with DNA-VLPs was due to TD, or T cell dependent, recall of B cell memory.

To contrast our findings on DNA-VLPs with protein-based materials, we employed ferritin-based P-VLPs with 24-valent display of RBDs (**P-VLP-24x**) on a scaffold of approximately 12 nm scaffold diameter (**Figure S1**)^32,39^. Following the validation of efficient B cell activation in vitro (**Figure S7**), C57BL/6 mice were sequentially immunized intraperitoneally with monomeric wild-type RBD, **I52-30x**, or **P-VLP-24x** at doses equivalent to 7.5 μg RBD (**Figure 4A**). While RBD-specific IgG titers were enhanced for both **I52-30x** and **P-VLP-24x** (**Figure 4B and 4C**), **P-VLP-24x** also exhibited boosted IgG titers against the ferritin protein scaffold itself (**Figure 4B**). In contrast, we did not observe boosting of DNA-specific IgG titers following immunization with DNA-VLPs, indicating an absence of B cell memory of the DNA-based scaffold (**Figure 4C and S8**). Importantly, this same behavior was observed when a higher DNA dose was administered (**I52-6x**) and also when monomeric RBD was administered (**Figure S8**).

**Figure 4.**
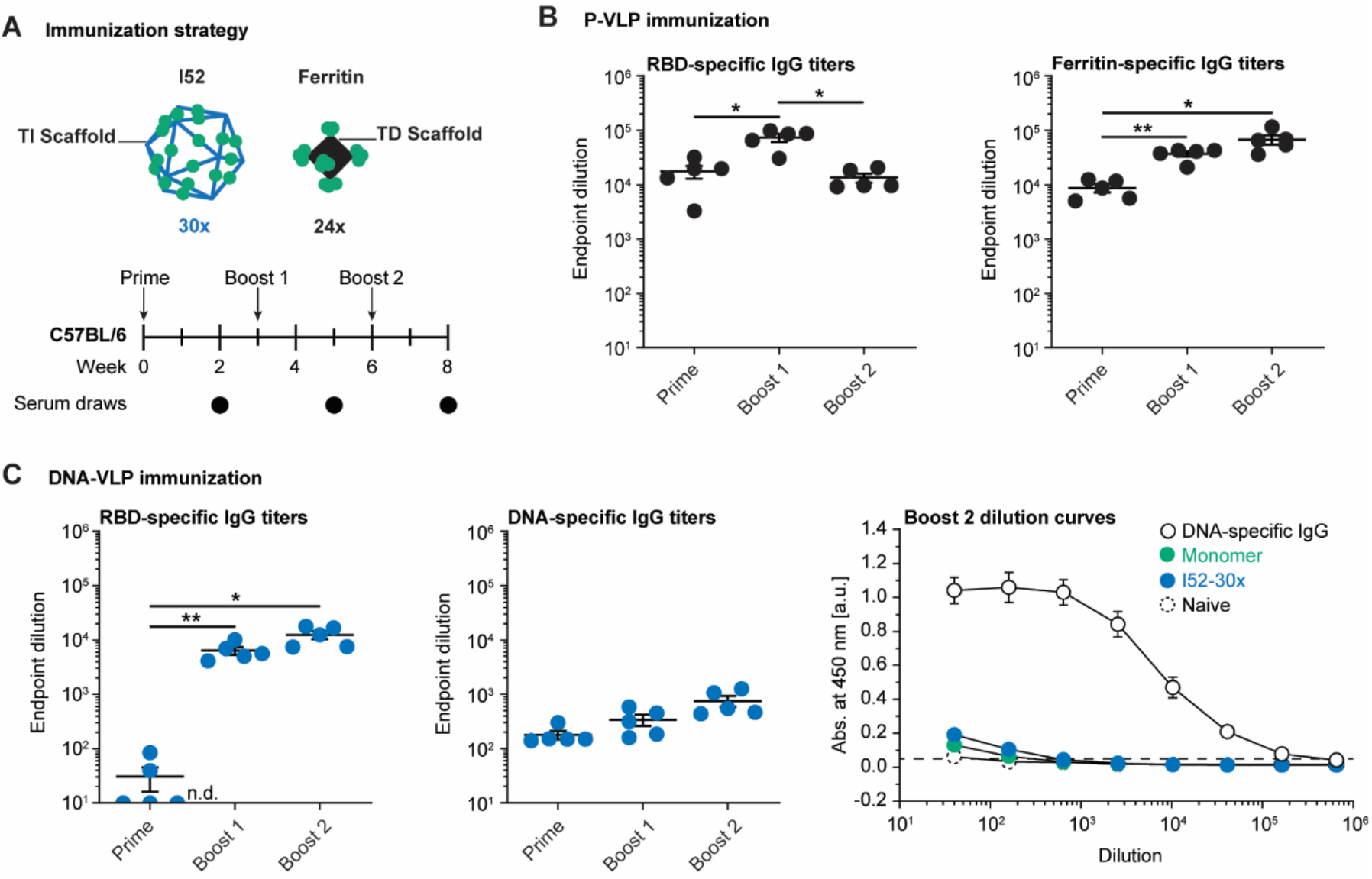
Scaffold-specific antibody responses to functionalized DNA-VLPs and P-VLPs. (A)Mice were sequentially immunized with monomeric RBD, RBD-functionalized DNA-VLP, or ferritin-based P-VLP. (B)RBD-specific and scaffold-specific IgG endpoint dilutions for the P-VLP immunization. **(C)** RBD-specific and scaffold-specific IgG endpoint dilutions for the DNA-VLP immunization; dilution curves for Boost 2 of the DNA-specific ELISA. The DNA-specific IgG control was diluted from 10 μg/ml. N=5 animals were used in each experimental group. One-way ANOVA was performed followed by Dunnett’s T3 multiple comparison test at α = 0.05. Error bars represent the standard error of the mean. Non-responder mice (denoted as n.d. = not detectable) were not considered for statistical analysis.

## Conclusions

Rational vaccine design seeks to leverage multivalent antigen display together with epitope-focusing to generate potent, neutralizing antibody responses^79^. P-VLPs have demonstrated limited control over antigen valency and antigen type^8,9,29,38^ together with low-cost, scalable production^27,30,32,38^, but they suffer from the generation of distracting, off-target antibody responses against the protein scaffold material itself^39,40,50^.

As an alternative VLP scaffold material, here we investigated the multivalent display of the RBD antigen from SARS-CoV-2 using icosahedral DNA origami VLPs. Our results suggest that DNA-VLPs covalently functionalized with 30 copies of the RBD antigen significantly enhance neutralizing antibody responses compared with monomeric RBD antigen alone, consistent with findings for P-VLPs displaying RBD^10,30,32,33,38,39^, as well as other antigens^8,9,23^. However, unlike P-VLPs that elicit scaffold-directed humoral immunity within the memory compartment^32,33,38,39^, compatible with our findings here for a ferritin-based P-VLP, DNA-VLPs did not generate B cell memory to the VLP scaffold material itself. Importantly, this suggests that the distraction of antibody responses away from the target protein antigen of interest in P-VLPs, which may reduce cross-neutralization of variants in the case of SARS-CoV-2^39^ and potentially result in irreversible off-target immune imprinting^43,44,45,80^, may be avoided by DNA-VLPs. While our finding was expected for TI antigens such as DNA, it is also well established that TD antibody responses can be generated for TI antigens covalently attached to protein scaffolds^81,82^. Our results, however, indicate that the inverse is not necessarily the default, namely scaffolding TD antigens with TI antigens does not generate boostable B cell memory against the TI scaffold for our vaccine platform. At the same time, we observed robust valency-dependent TD antibody responses to the RBD, akin to virosomal and ISCOM-based vaccine designs in which protein antigens are multivalently displayed by TI antigen-composed matrices^83-86^.

Because DNA origami uniquely offers simultaneous yet independent control over spatial antigen display, scaffold size, and scaffold geometry, we were able to investigate the impact of antigen valency alone on antibody responses using an optimally sized, geometrically fixed ∼35 nm icosahedral VLP scaffold, whereby 30 but not six copies of RBD were found to be sufficient to enhance neutralizing antibody titers compared with monomeric RBD alone. Future work may seek to additionally examine the impact of antigen spacing with fixed valency^58^, as well as alternative scaffold sizes and geometries, which might ultimately be required to resolve critical thresholds for enhancing antibody responses beyond monomeric antigens, as well as to optimize B cell responses for certain antigens^9,29,30^. Additional important extensions to our study include examining GC formation in B cell follicles, which is known to be important to generate broadly neutralizing antibodies and long-lived humoral immunity^87-89^. Toward this end, it will be interesting to investigate to what extent multivalent antigen display by DNA-VLPs is maintained in secondary lymphoid organs, particularly in the presence of endonuclease^90,91^ and protease^89^ degradation; what the breadth of neutralizing antibody responses is; and what the longevity of humoral immunity is. To further enhance B cell activation and trafficking to secondary lymphoid organs, DNA-VLP valency may be increased, slow escalating dosing may be used^92^, and active follicle targeting with carbohydrates may be incorporated^5,87^. Co-formulation with TLR-based adjuvants may also be used to enhance T cell help, drive GC reactions, and improve humoral immunity^93-95^. Finally, DNA stabilization strategies^91^ may be needed to increase the longevity of DNA-VLPs within follicles and GCs in the presence of endonucleases.

Beyond rational vaccine design and promoting antibody focusing, our discovery that DNA scaffolds are not neutralized by DNA-specific antibodies is of significant importance towards enabling redosing in therapeutic nucleic acid delivery by avoiding antibody-dependent clearance^46,47^, whereby DNA origami may offer an important delivery vector^96^.

## Methods

Methods are described in the **Supporting Information**.

## Acknowledgments

E.-C.W., G.K., and M.B. were supported by NIH R21-EB026008, NSF CCF-1564025, CBET-1729397, and CCF-1956054, ONR N00014-21-1-4013 and N00014-17-1-2609, ARO ISN W911NF-13-D-0001, and Fast Grants Agreement EFF 4/15/20. E.-C. W. was additionally supported by the Feodor Lynen Fellowship of the Alexander von Humboldt Foundation. B.M.H was supported by NIGMS (T32GM007753) and F30 AI160908. J.F. was supported by NIGMS (T32AI007245). A.G.S. was supported by NIH (R01 AI146779) and a Massachusetts Consortium on Pathogenesis Readiness (MassCPR) grant. D.L was supported by NIH (R01AI124378, R01AI153098, R01AI155447, U19AI057229, P30AI060354). The SARS-CoV-2 virus work was performed at the Ragon Institute’s Biosafety Level 3 facility, which is supported by the NIH-funded Harvard University Center for AIDS Research (P30 AI060354) and the Massachusetts Consortium on Pathogen Readiness (MassCPR).

## Data availability

The DNA sequences used to assemble the wireframe DNA origami are included in Tables S1-3. All data files used to generate the figures in this manuscript are available from A. S., D. L. or M. B. upon request.

## Software and code availability

No previously unreported code or software were used for data collection and analysis.

