## Supplementary material for "Enhancing antibody responses by multivalent antigen display on thymus-independent DNA origami scaffolds": Methods, Supporting Notes, Supporting Figures and Supporting Tables

#### Title

### Methods

#### Materials

SS320 E. coli cells were purchased from Lucigen. EndoFree GigaPrep Kits were purchased from Qiagen. Tris acetate-EDTA (TAE) and PBS buffer were purchased from Corning. Oligonucleotide staples (purified via desalting columns) were ordered from IDT. The standard base and dibenzocyclooctyne-triethylene glycol (DBCO-TEG) phosphoramidites as well as ancillary chemicals and desalting columns were purchased from Glen Research. Organic solvents were purchased from Sigma Aldrich and VWR International. Agarose was purchased from IBI Scientific. The DNA agarose gel electrophoresis (AGE) standard (Quick-Load Purple 2-Log DNA ladder 0.1-10 kb) was purchased from New England Biolabs. $MgCl_2$, NaCl, $PEG_{8kDa}$, antibiotics, 2x YT medium, TE buffer, ethidium bromide (EtBr), TritonX-144 was purchased from Sigma-Aldrich. Amicon Ultra centrifugal filters (10 kDa and 100 kDa) and dialysis membranes (mixed cellulose ester, 0.025 µm) were purchased from Sigma Aldrich. Zeba spin columns (7 kDa) and Proteinase K were purchased from ThermoFisher Scientific. Transmission electron microscopy grids (CF200H-CU) were obtained from Electron Microscopy Sciences. Uranyl formate was purchased from Sigma Aldrich. The codon-optimized gene for the expression of the SARS-CoV-2 receptor binding domain (RBD) was purchased form IDT. QuickChange Mutagenesis Kits were purchased from Agilent. Expi293 Expression System Kits were purchased from ThermoFisher Scientific. TALON cobalt resin was purchased from Takara. Superdex 200 Increase columns were purchased from Cytiva. Ellman's reagent was purchased from Sigma Aldrich. Amino-$PEG_3$-azide was purchased from BroadPharm. SMCC was purchased from ThermoFisher Scientific. ToxinSensor Gel Clot Endotoxin Assay Kits were purchased from GenScript. 96- and 384-well cell culture plates were purchased from Corning and Greiner, respectively. 96-well MaxiSorp plates were purchased from Nunc. Goat anti-human-Fc-PE antibody conjugates were purchased from SouthernBiotech. Mouse anti-human-IgM-APC antibody conjugates were purchased from BioLegend. Mouse anti-human-κ-light-chain-PE antibody conjugates were purchased from eBioscience. RPMI 1640 was purchased from Sigma Aldrich. Cy5-Az and Cy5-DBCO were purchased from Sigma Aldrich. Calf-thymus DNA was purchased from Sigma Aldrich. Mouse anti-dsDNA antibodies were purchased from Abcam. 1% casein blocking buffer was purchased from G-Biosciences. Goat anti-human-IgM was purchased from Jackson ImmunoResearch. 100x penicillin-streptomycin-glutamine solution was purchased from ThermoFisher Scientific. For neutralization assays, Vero-E6 (Cat#CRL-1586, ATCC) and A549-hAce2 (Cat# NR-53821, NEI resources) cell lines were maintained in D10+ media (DMEM (Corning) supplemented with HEPES (Corning), 1X Penicillin 100 IU/mL/Streptomycin 100 µg/mL (Corning), 1X Glutamine (Glutamax, ThermoFisher Scientific), and 10% Fetal Bovine serum (FBS, Sigma)) at 37C and 5% CO2. SARS-CoV-2 USA-WA1/2020 viral stock was expanded from BEI Resources reagent NR52281 on Vero-E6 cells and quantified by plaque assay.

#### Scaffold synthesis

The custom-length DNA scaffold (phI52, 3210 nt) for **I52** was prepared as previously described **(Table S1)**[1]. Briefly, SS320 E coli cells were transformed with the phI52 plasmid, based on the pUC19 vector, and the M13cp helper plasmid (generously provided by Andrew Bradbury, Los Alamos National Laboratories). Next, pre-cultures of the transformed cells were grown overnight at 37°C, diluted 100-fold and incubated for another 8 h, with all steps using 2x YT medium containing 100 µg/mL ampicillin, 15 µg/mL chloramphenicol and 5 µg/mL of tetracycline. Cells were sedimented by centrifuging three times at 4000 g for 3 min and subsequently discarded. Phage was precipitated from the supernatant in presence of 6% (w/v) $PEG_{8kDa}$ and 3% (w/v) of NaCl by stirring at 4°C for 1 hour and harvested by centrifugation at 20,000 g at 4°C for 1 h. After resuspension in TE buffer, ssDNA was extracted via the EndoFree GigaPrep purification protocol with the following modifications: Proteinase K was added to buffer P1 followed by incubation at 37°C for 1 h, addition of buffer P2 and incubation at 70°C for 10 minutes. After ssDNA purification, Triton X-144 was used to remove residual endotoxins to levels less than 0.2 EU/µmol DNA scaffold, corresponding to less than 0.000015 EU/injection into mice[2]. Endotoxin levels were measured using ToxinSensor Gel Clot Endotoxin Assay Kits. Purity of the scaffold was analyzed by agarose gel electrophoresis (AGE) (1.6% agarose, TAE buffer with 12 mM $MgCl_2$, EtBr, 65V for 150 in at 4°C).

**Oligonucleotide staple synthesis**

Solid-phase DNA synthesis was performed on a Dr. Oligo synthesizer purchased from Biolytic. DNA synthesis was performed on a 200 nmol scale, starting from universal 1000 Å CPG solid-supports and following the standard protocol[3]. Standard base and DBCO-TEG phosphoramidites were dissolved in anhydrous acetonitrile to afford 0.1 M solutions and were used in 10-fold excess at a coupling time of 10 min. Coupling efficiency was monitored after removal of the dimethoxy trityl (DMT) 5'-OH protecting groups. After solid-phase synthesis, oligonucleotides were cleaved off the resin in concentrated ammonium hydroxide at 60°C for 3 h and purified over desalting columns. Installation of the DBCO-TEG modification was characterized by reversed-phase high-performance liquid (HPLC) and modified staples were purified as previously described[4]. For the assembly of **I52-5x-Cy5** and **I52-30x-DBCO-5x-Cy5**, Cy5-modified oligonucleotide staples were synthesized preassembly. 10x excess of Az-Cy5 was added to 50 µM DBCO-modified oligonucleotide staple in PBS with 10% DMF and incubated overnight at room temperature. Excess dye was removed using NAP-5 columns prior to purification of Cy5-modified oligonucleotide staples via reversed-phase HPLC as previously described[4].

**Antigen synthesis**

The codon-optimized gene for the expression of the RBD of SARS-CoV-2 (GenBank ID = MN975262.1; residues 319-529) was cloned into a pVRC vector containing a C-terminal HRV C3 protease cleavage site followed by 8x His and SBP tags. An additional C-terminal Cys residue was inserted using QuickChange Mutagenesis following the manufacturer's protocol and mutagenesis was confirmed by next-generation sequencing (Azenta) to afford RBD-Cys. HEK Expi293F cells were transiently transfected with the RBD plasmid using Expifectamine following the manufacturer's protocol. After 5 to 7 days, supernatants were harvested by centrifugation at 4000 g at room temperature for 5 min and the RBD-Cys was purified into PBS by affinity chromatography using TALON cobalt resin followed by size-exclusion chromatography using Superdex 200 Increase columns and stored at 4°C for less than 7 days.

Antigen modification with the azide linker was adapted from published protocols[4]. Briefly, 100 µM RBD-Cys was incubated with 10x excess of TCEP in PBS for 30 minutes at room temperature and subsequently purified into PBS with 10 mM EDTA using Zeba spin columns (7 kDa). Incubation for approximately 6 h at room temperature allowed for the reoxidation of disulfides prior to the addition of the azide linker as monitored by Ellman's assay. The azide linker was assembled by incubation of 60 mM SMCC with 1.1x excess of amino-$PEG_3$-azide for 1 h at room temperature. Subsequently, 10x excess of crude azide linker was added to reduced antigen and the reaction mixture was incubated overnight at room temperature to afford RBD-Az. RBD-Az was purified into PBS using Amicon Ultra centrifugal filters (10 kDa, 5000 g) followed by size-exclusion chromatography using Superdex 200 Increase columns and stored at 4°C for less than 7 days. To the label azide-modified antigen with dyes, 50 µM RDB-Az was incubated with 5x excess of DBCO-Cy5 in PBS for 30 min at room temperature and subsequently purified into PBS using Amicon Ultra centrifugal filters (10 kDa, 5000 g). RBD concentrations and dye labeling efficiency were determined by absorbance measurments at 280 nm (ε = 39400 1/(M·cm) and M = 31 kDa) and 650 nm (ε = 250000 1/(M·cm)).

**DNA origami design and assembly**

**I52**, **I52-1x-DBCO**, **I52-6x-DBCO** and **I52-30x-DBCO** as well as **I52-5x-Cy5** and **I52-30x-DBCO-5x-Cy5** were designed using DAEDALUS and nick position of edge staples were adjusted manually for outward orientation **(Table S2 and S3)**[5]. The DNA virus-like particles (DNA-VLPs) were assembled as previously described[5]. Briefly, 30 nM of scaffold and 300 nM of each oligonucleotide staple were dissolved in TAE buffer with 12 mM $MgCl_2$ and thermally annealed as follows: 95°C for 5 min, 80–75°C at 1°C per 5 min, 75–30°C at 1°C per 15 min, and 30–25°C at 1°C per 10 min. The DNA-VLPs were purified into PBS using Amicon Ultra centrifugal filters (100 kDa, 2000 g) and stored at 4°C. Purity and monodispersity of the DNA-VLPs were validated by AGE (1.6% agarose, TAE buffer with 12 mM $MgCl_2$, EtBr, 65V for 150 min at 4°C) **(Figure S1)**.

**DNA origami functionalization**

DBCO-bearing DNA-VLPs were functionalized with RBD-Az to yield **I52-1x**, **I52-6x**, **I52-30x** or **I52-Cy5-30x**. At least 150 nM of **I52-30x-DBCO** or **I52-30x-DBCO-5x-Cy5** and at least 750 nM of **I52-1x-DBCO** or **I52-6x-DBCO** were incubated with 30x excess per DBCO group of RBD-Az in PBS at room temperature for 24 h. RBD-functionalized DNA-VLPs were purified into PBS by drop dialysis (mixed cellulose ester, 0.025 µm) and stored at room temperature for less than 7 days. Purity and monodispersity of the DNA-VLPs were validated by AGE (1.6% agarose, TAE buffer with 12 mM $MgCl_2$, EtBr, 65V for 150 in at 4°C) and dynamic light scattering (DLS, Malvern Zetasizer Ultra). Coverage with antigens was measuring using tryptophan fluorescence spectroscopy at 100 nM RBD and denaturing, reversed-phase HPLC as previously described[4].

**Transmission electron microscopy**

Uranyl formate staining of DNA-VLP samples was adapted from an existing protocol[6]. Briefly, **I52-30x** was diluted to 5 nM, and 5 ul of the solution were immediately deposited onto glow-discharged electron microscopy grids. After 30s, the solution was removed by blotting with filter paper and the grids were washed with 5 ul of freshly prepared 2% uranyl formate with 5 mM NaOH. After removal of the washing solution by blotting, 15 ul of the uranyl formate solution was added, incubated for 30 s and the removed by blotting. Finally, the grids were dried in vacuo and transmission electron microscopy (TEM, FEI Tecnai G2 Spirit Twin) was conducted at 120 keV.

**Protein-based virus-like particle synthesis**

The SARS-CoV-2 RBD with C-terminal 8x His and SpyTags was expressed and purified as described above. H. pylori ferritin nanoparticles were expressed with N-terminal SpyCatcher and 8x His tags were expressed and purified as previously described[7]. SpyTag-SpyCatcher conjugation was performed overnight at 4°C at 4x excess of RBD-SpyTag per SpyCatcher. 24-

valent P-VLPs bearing RBD (**P-VLP-24x**) were subsequently purified into PBS by size exclusion chromatography as previously described[7].

**Biolayer interferometry**

Biolayer interferometry binding experiments were performed after immobilization of CR3022 and B38 Fab at 0.1 mg/ml on FAB2G sensors (Sartorius, BLItz). Wild-type RBD and RBD-Az served as analytes at 10 µM in the manufacturer's Kinetics Buffer.

**ACE2-expressing cell binding assay**

ACE-2 expressing HEK 293T cells (generously provided by Nir Hacohen and Michael Farzan, Massachusetts General Hospital and The Scripps Research Institute) were harvested and washed with PBS with 2% FBS. 200000 cells per well were transferred to 96-well cell culture plates and 100 ul of wild-type RBD, RBD-Cy5 and **I52-30x-RBD-5x-Cy5** at concentration corresponding to 200 nM RBD in PBS were added. The concentration of **I52-5x-Cy5** corresponded to that of **I52-30x-RBD-5x-Cy5**. Following incubation for 60 min on ice, cells were washed twice with PBS with 2% FBS and stained with 50 ul at 200 nM of the anti-RBD antibody CR3022 for 30 min at room temperature, following pre-complexed with goat anti-human-PE at 200x excess. The suspension was protected from light and incubated for 30 min on ice, washed twice with PBS with 2% FBS and resuspended in 100 ul PBS with 2% FBS. For RBD-Cy5, **I52-5x-Cy5** and **I52-30x-RBD-5x-Cy5** cell binding was also detected via Cy5 fluorescence. After incubation with RBD or DNA-VLPs, no staining was performed and cells were directly resuspended in 100 ul PBS with 2% FBS. Cell binding was analyzed by flow cytometry (Strategim S1000Exi Flow Cytometer) and data processing was conducted using FlowJo (BD Biosciences, v.10). Cy5 fluorescence intensities obtained by flow cytometry were corrected according the relative brightness of RBD-Cy5, **I52-5x-Cy5** an **I52-30x-RBD-5x-Cy5** as quantified by fluorescence spectroscopy in PBS.

**B-cell activation assay**

The B-cell activation assay was adapted from previously established protocols[8-10]. Briefly, the human anti-RBD antibodies CR3022[11] and B38[12] were expressed as IgM B-cell receptors (BCRs) in Ramos B cells after lentiviral transfection of the corresponding light chain and transmembrane IgM heavy chain genes. 5 to 7 days after transfection, BCR-expressing Ramos B cells were FACS-sorted for IgM and κ light chain expression (Sony Biotech SH800S Cell Sorter). Sorted cells were expanded in RPMI supplemented with 15% FBS and 1x penicillin-streptomycin-glutamine and 1,000,000 cells were harvested and resuspended RPMI with Fura Red solution following the manufacturer's protocol. After incubation for 30 min at 37°C, cells were harvested and resuspended in 2 ml RPMI medium prior to detection of BCR signaling by flow cytometry at 637 nm. Following 30 s of baseline data acquisition, wild-type RBD, **I52-1x-RBD**, **I52-6x-RBD**, **I52-30x-RBD** or **P-VLP-24x-RBD** at concentrations corresponding to 30 nM RBD were added before continuing data acquisition for an additional 270 s. The concentration of **I52** corresponded to that of **I52-30x-RBD**. Goat anti-human IgM at 10 µg/ml served as a positive control. Maximum total $Ca^{2+}$ flux was measured after addition of 10 µg/ml ionomycin. Fluorescence traces were processed as follows: For each trace, the average fluorescence of the 30 s baseline data acquisition was subtracted. Next, the fluorescence traces were normalized to the $Ca^{2+}$ flux induced by ionomycin to obtain relative $Ca^{2+}$ flux traces. The total $Ca^{2+}$ flux was quantified by integration to obtain normalized areas under the curve (AUC).

**Immunization experiments**

Wildtype C57BL/6 or *Tcra*$^{-/-}$ mice[13] (n = 5 per group, males and females, 6-8 weeks of age) were pre-bled and then sequentially immunized intraperitoneally with wild-type RBD, **I52-1x-RBD**, **I52-6x-RBD**, **I52-30x-RBD** or **P-VLP-24x-RBD** at equimolar doses equivalent to 7.5 µg RBD. The antigens were injected in 100ul containing 50% Sigma adjuvant as performed previously [10, 14, 15]. Immunization occurred at weeks 0, 3 and 6 and blood draws occurred two weeks after each immunization.

**RBD- and ferritin-specific IgG ELISA**

Recombinant RBD (or in some cases naked ferritin) was used to coat MaxiSorp plates at 200 ng/well overnight at 4°C. The plates were washed with PBS, blocked with PBS with 3% non-fat milk for 1 h at room temperature and subsequently washed with PBS. Mouse sera or human anti-RBD antibodies were diluted in PBS, transferred to the plates and incubated for 1 h at room temperature. The plates were washed with PBS with 0.05% Tween-20 and subsequently incubated with a 1/5000 dilution of either goat anti-mouse-IgM (IgG-HRP, GE Healthcare) or sheep anti-murine IgG (HRP-IgG, GE Healthcare). Following washing with PBS with 0.05% Tween-20, TMB was added, and the developer reaction was then stopped using 1N sulfuric acid. The plates were read by absorbance at 450 nm (Tecan Infinite m1000 Pro microplate absorbance reader (Mannedorf, Switzerland). Antibody endpoint dilutions were interpolated using were calculated using an absorbance cut-off of 0.05 (Graphpad Prism version 9.1, GraphPad Software Inc). The loading of RBD was also standardized to recombinant mAbs CR3022 and B38, which we expressed and purified previously (PMIDs: 36423634, 35303475, 33594359). In this case ELISA was as above, except that sheep anti-human IgG-HRP (GE Healthcare) was used as the secondary antibody.

**DNA-specific IgG ELISA**

Calf-thymus DNA was reconstituted in water 50 µg/ml and used to coat 96-well MaxiSorp plates overnight at 4°C. The plates were washed with PBS, blocked with 1% casein buffer for 2 h at room temperature and subsequently washed with PBS. Mouse sera or mouse anti-dsDNA antibody were diluted in blocking buffer, transferred to the plates and incubated for 2 h at room temperature. The plates were washed with PBS with 0.2% Tween-20 and incubated with 1:5000 dilution XX dilution goat anti-mouse-IgG-HRP antibody (BioRad) for 1 h at room temperature. Following washing with PBS with 0.2% Tween-20, TMB was added, the plates were incubated for 3 min and the reaction was stopped using sulfuric acid. DNA-specific IgG titers were determined by absorbance measurements at 450 nm. Endpoint dilutions were calculated using an absorbance cut-off of 0.05 as determined from the limit of detection determined for PBS-incubated wells Graphpad Prism version 9.1, GraphPad Software Inc).

**Pseudovirus neutralization assay**

SARS-CoV-2 neutralization was assessed using pseudotyped lentivirus particles expressing S glycoprotein trimer as previously described[16]. Briefly, pseudovirus corresponding to the wild-type, Wuhan strain were produced by transient transfection of HEK 293T cells. The titers of viral supernatants were determined via flow cytometry with ACE2-expressing HEK 293T cells and via the HIV-1 p24$^{CA}$ antigen capture assay (Leidos Biomedical Research) Assays were performed in 384-well plates using a fluorescence plate reader (Tecan Fluent Automated Workstation). Mouse sera (or CR3022 and B38 mAb standards), starting from a 3x initial dilution, were serially 3x diluted in 20 µl followed by the addition of 20 µl of pseudovirus containing 250 ifu and incubated for 1 h at room temperature. Next 1,00,000 ACE2-expressing HEK 293T cells are were added per well and incubated for 60 to 72 h at 37°C. After transfection, cells were lysed and incubated on a shaker for 5 min at room temperature before measuring luciferase expression (Molecular Devices SpectraMax L). Relative neutralization for each serum dilution was calculated after subtracting background luminescence and diving by the luminescence in absence of sera. $NT_{50}$ vales were derived by fitting the Dose-Response equations to serum dilution curves.

**Native virus neutralization assay**

SARS-CoV-2 neutralization was additionally assessed using live virus as previously described[17]. Briefly, A549-hAce2 cells were detached using Trypsin-EDTA (Fisher Scientific) and seeded at 40,000 cells per well in 96-well plates 16-20 hours before infection. Four hours before infection, the cell culture supernatant was removed and 75uL of D2+ media was added (2% FBS instead of 10%).

Mice sera was diluted in D+ media (no FBS) in 3-fold serial dilutions, mixed 1:1 (v/v) with SARS-CoV-2 diluted at 40,000pfu/mL and incubated at 37C and 5% CO2 for 1 hr. Twenty-five microlitres of the sera-virus solution was added in triplicate wells for each condition for a final multiplicity of infection of 0.01 (virus to cell). The 96w plates were centrifuged for 30min at 2,000x g at 37C and then incubated at 37C and 5% CO2 for 48h. Each plate included a no infection control, a no treatment control maximum infection as well serially diluted B38 mAb as a positive control.

After 48 hours, the cell culture supernatant was discarded, the cells were washed with PBS (Corning) then harvested using TrypLE (Life Technologies) and flow cytometry buffer (2% FBS in PBS). Cells were washed, then stained with live/dead fixable blue stain (Thermo Fisher) for 30 minutes at 4°C. After one wash with flow cytometry buffer, the cells were fixed using 4% paraformaldehyde (Santa Cruz) for 30 minutes at 4°C. The fixed cells were removed from the BSL3 laboratory and prepared for intracellular staining using Perm/Wash buffer (BD Biosciences). The permeabilized cells were stained with mouse anti-SARS-CoV-2 Nucleocapsid antibody (Biolegend) for 30 minutes at 4°C, then washed and stained with secondary antibody labelled with Pe-Cy7 (Biolegend) for 30 minutes at 4°C. Finally, the cells were washed and resuspended in flow cytometry buffer.

Flow cytometry was performed on a BD Symphony (BD Biosciences). FCS files were analyzed using FlowJo software (version 10). Additional data analysis was performed using GraphPad Prism (version 9) to fit curves for the neutralization data.

#### Supporting Notes

##### Note S1 – Fabrication of RBD-functionalized DNA-VLPs

X-ray structures of the ACE2-RBD complex suggested that modifications at the C-terminus of the antigen would allow for recognition of neutralizing epitopes including the receptor binding motif (RBM)[18]. We cloned and expressed recombinant RBD bearing a C-terminal His-tag followed by an additional Cys using HEK Expi293F cells (RBD-Cys, **Figure S2**). Incubation of RBD-Cys with TCEP revealed that at least 1 of the 4 disulfide bridges was susceptible to reduction as determined by Ellman's assay (**Figure S2**). To ensure selective labeling of the C-terminal Cys, the reduced antigen was purified into PBS containing 10 mM EDTA and incubated at room temperature to allow for reoxidation. After approximately 6 h, the number of reduced Cys stabilized at 1 per RBD, indicating kinetically favored reoxidation of the disulfide bridges over the C-terminal Cys (**Figure S2**). The reduced RBD-Cys was reacted with an SMCC-TEG-azide linker to afford RBD-Az. Conversion of the linker reaction was validated by selective labeling of the azide with Cy5-DBCO and subsequent ratiometric absorbance measurements. In addition to flow cytometry experiments with ACE2-expressing HEK293 cells, the preservation of the RBM for RBD-Az was validated by biolayer interferometry. Reaction conditions for the *in situ* functionalization of DBCO-bearing **I52** were screened and set to at least 150 nM VLP at 30x excess of RBD-Az per DBCO to achieve near-quantitative coverage (**Figure S2**). Following 24 h incubation at room temperature, **I52-1x, 6x**,and **30x** were purified by drop dialysis. Notably, functionalization efficiency was dependent on maximum DBCO concentrations rather than nanoparticle concentrations, and we were only able to obtain 30% functionalization efficiency for **I52-1x**.

#### Supporting Figures

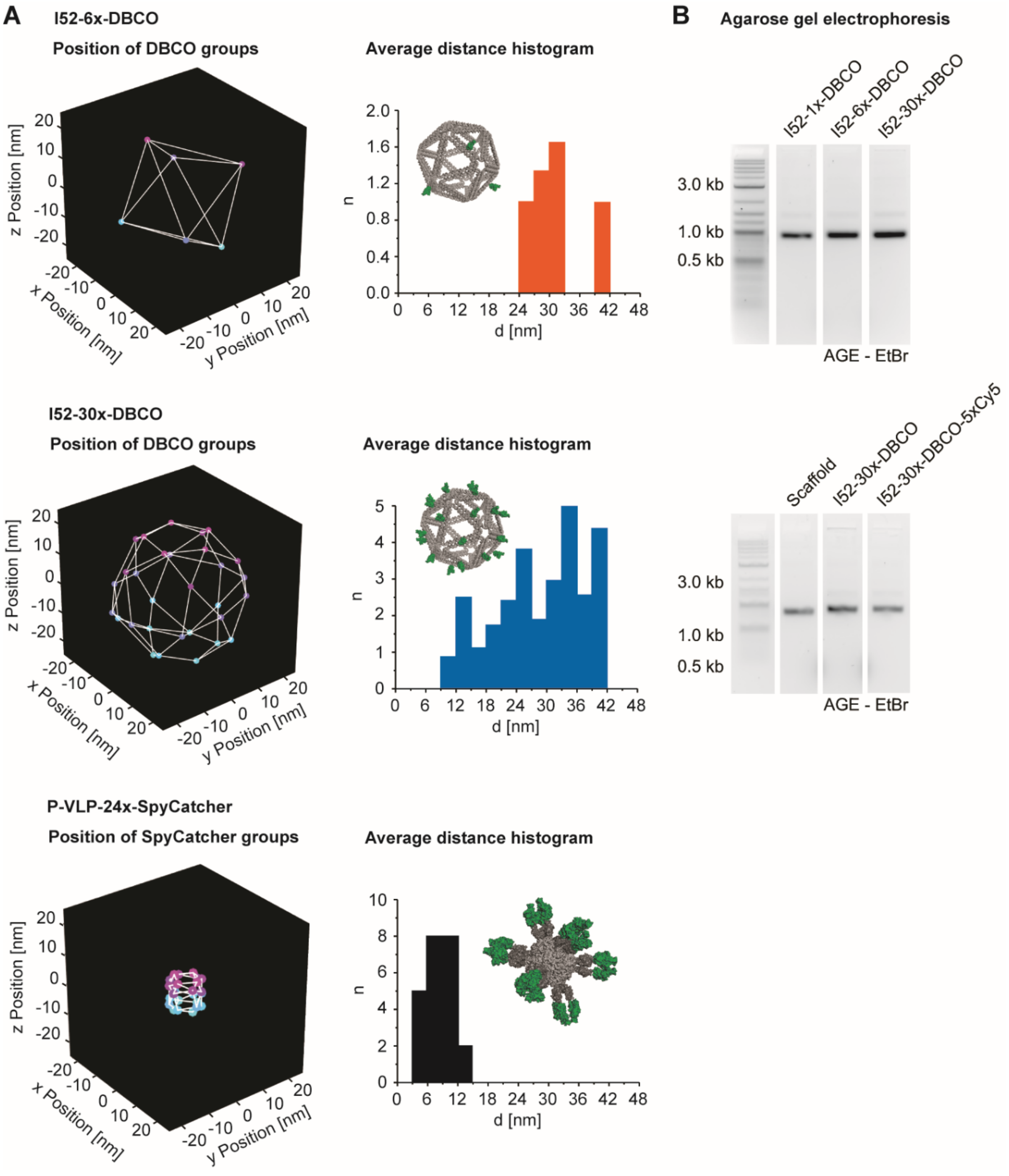


**Figure S1 – Supporting information for DNA-VLP and P-VLP assembly.**
**(A)** The positions of functional groups were mapped by proxy of the 5'OH groups of modified staples for **I52-6x-DBCO** and **I52-30x-DBCO** and of the SpyCatcher locations for **P-VLP-24x-SpyCatcher**. These mapped positions served to calculate distance histograms, using the average, rank-ordered distances between each functional group and all neighboring conjugation sites. **(B)** AGE served to validate the assembly of DBCO-functionalized DNA-VLPs.

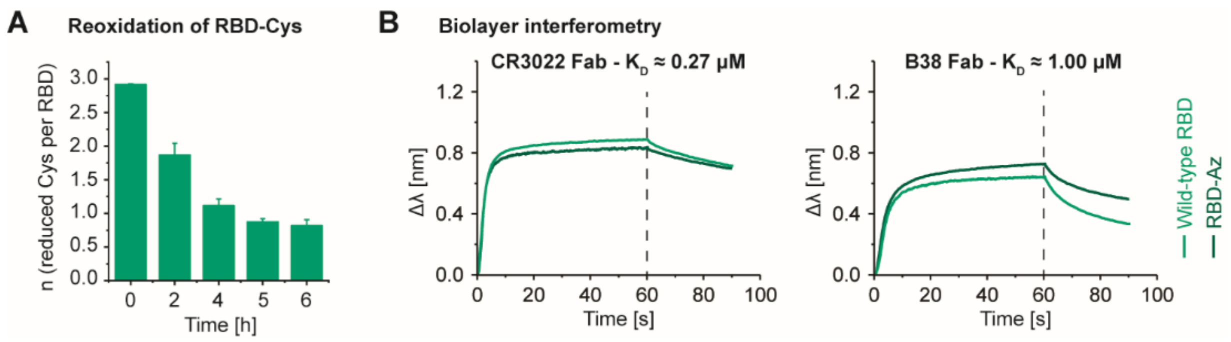


**Figure S2 – Supporting information for antigen expression and functionalization.**
**(A)**The reduction and subsequent reoxidation of RBD-Cys in PBS with 10 mM EDTA was monitored by Ellman's assay. The reoxidation kinetics revealed that at least 1 of the 4 disulfide bridges was susceptible to reduction but reoxidation was favored compared to the additional, C-terminal Cys, allowing for selective modification with an SMCC-TEG-azide linker. **(B)** In addition to flow cytometry experiments with ACE2-expressing HEK293 cells, the preservation of the RBM for RBD-Az was validated by biolayer interferometry at 10 µM antigen. Binding kinetics for both the CR3022 and B38 Fab fragments were comparable to wild-type RBD. All experiments were conducted at n = 3 biological replicates. Error bars represent the standard error of the mean. Representative biolayer interferometry curves are shown.

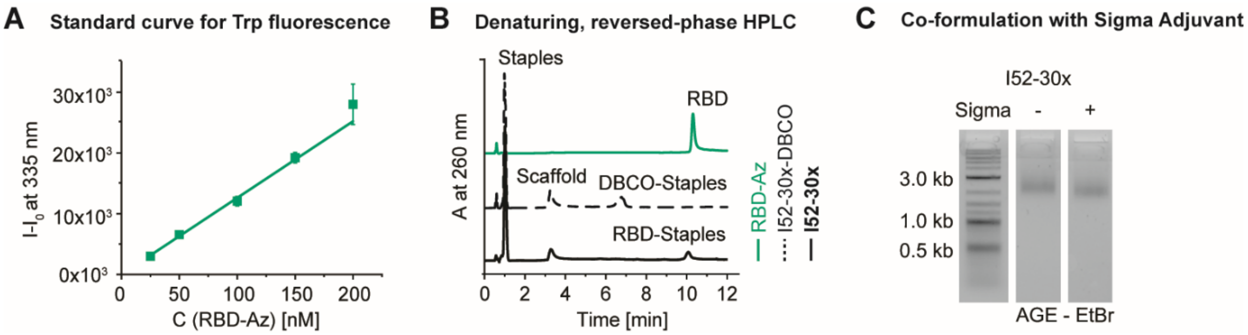


**Figure S3 – Supporting information for DNA origami functionalization.**
**(A)** RBD coverage of DNA-VLPs was determined by Trp fluorescence as previously described for other antigens[19]. The corresponding standard curve for RBD-Az is shown. **(B)** Near-quantitative reaction conversion was further validated for **I52-30x** via denaturing, reversed-phase HPLC[4]. **(C)** 100 nM **I52-30x** were co-formulated with PBS (left) or Sigma adjuvant (right) at equal volumes and characterized by AGE, supporting their compatibility. Trp fluorescence values were determined from n = 3 biological replicates.

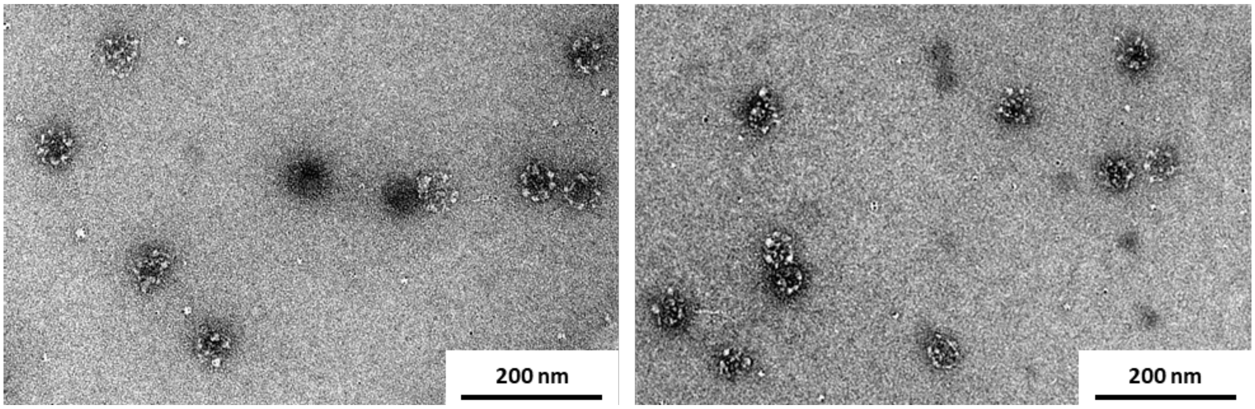


**Figure S4 – Supporting information for transmission electron microscopy.**
Representative TEM micrographs of **I52-30x** are shown. 2% uranyl formate was used for negative staining.

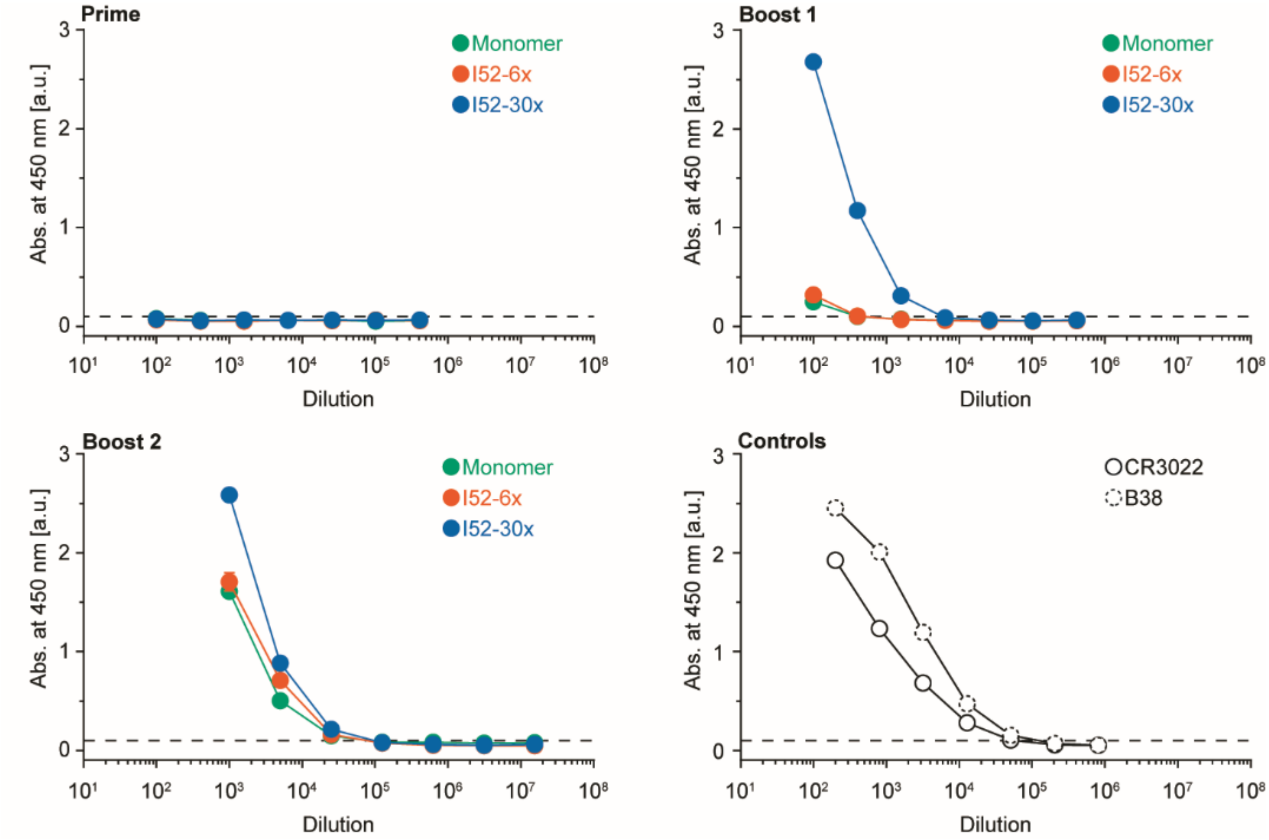


**Figure S5 – Supporting information for RBD-specific IgG ELISA.**
Dilution curves for RBD-specific IgG ELISAs with mouse sera following immunization with 7.5 µg RBD are shown. Plates were coated with C-terminally cleaved RBD, devoid of 8x His and SBP tags. Absorbance values were determined from n = 5 biological replicates. Error bars represent the standard error of the mean. Representative dilutions curves for RBD-specific control IgGs are shown.

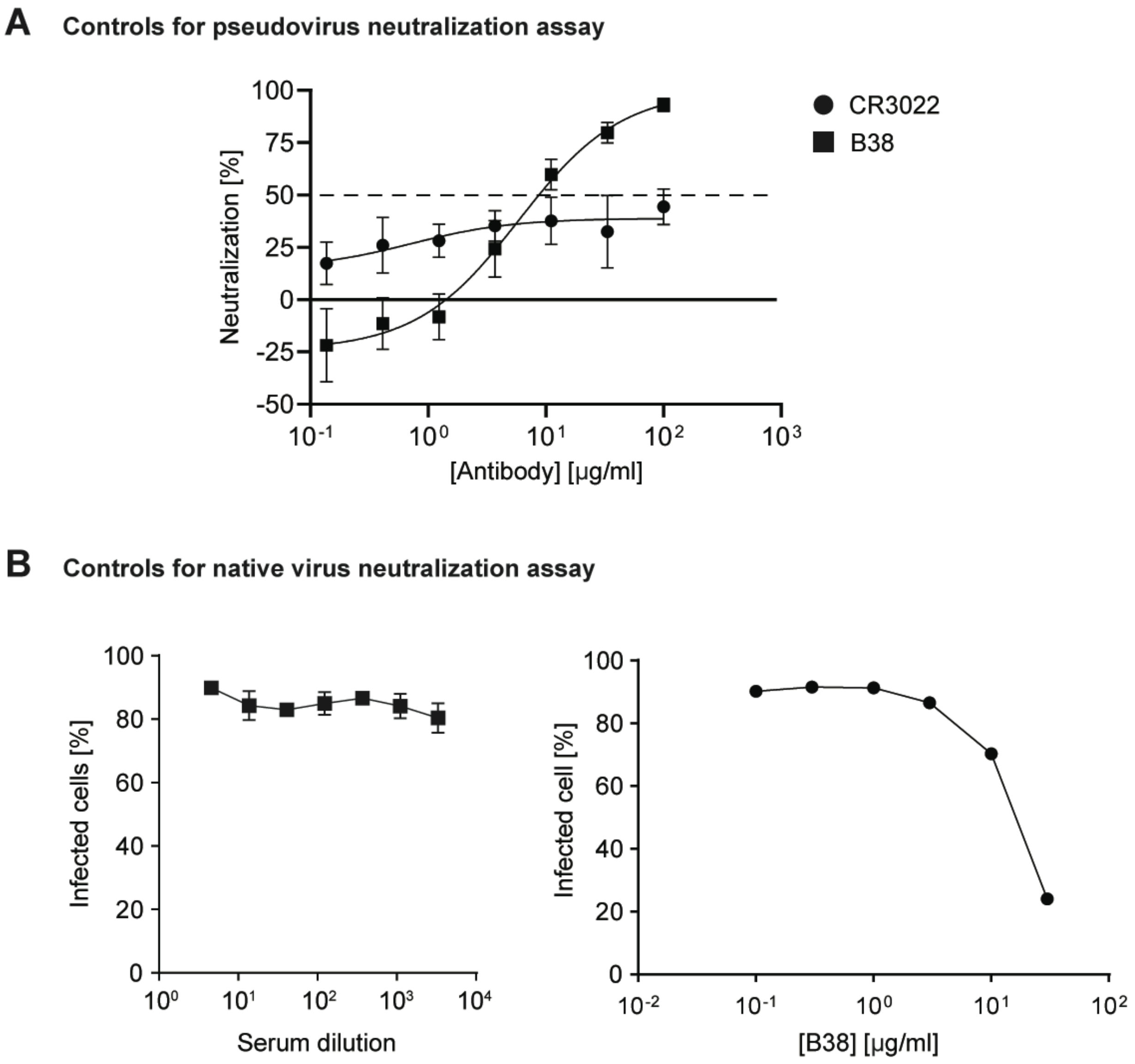


**Figure S6 – Supporting information for neutralization assays.**
**(A)** Dose-response curves for positive (B38) and negative (CR3022) control IgGs for the pseudovirus neutralization assay. **(B)** Dose-response curves for negative (serum from naïve mice) and positive (B38 IgG) controls in the native virus neutralization assay are shown. Error bars represent the standard error of the mean (n=5).

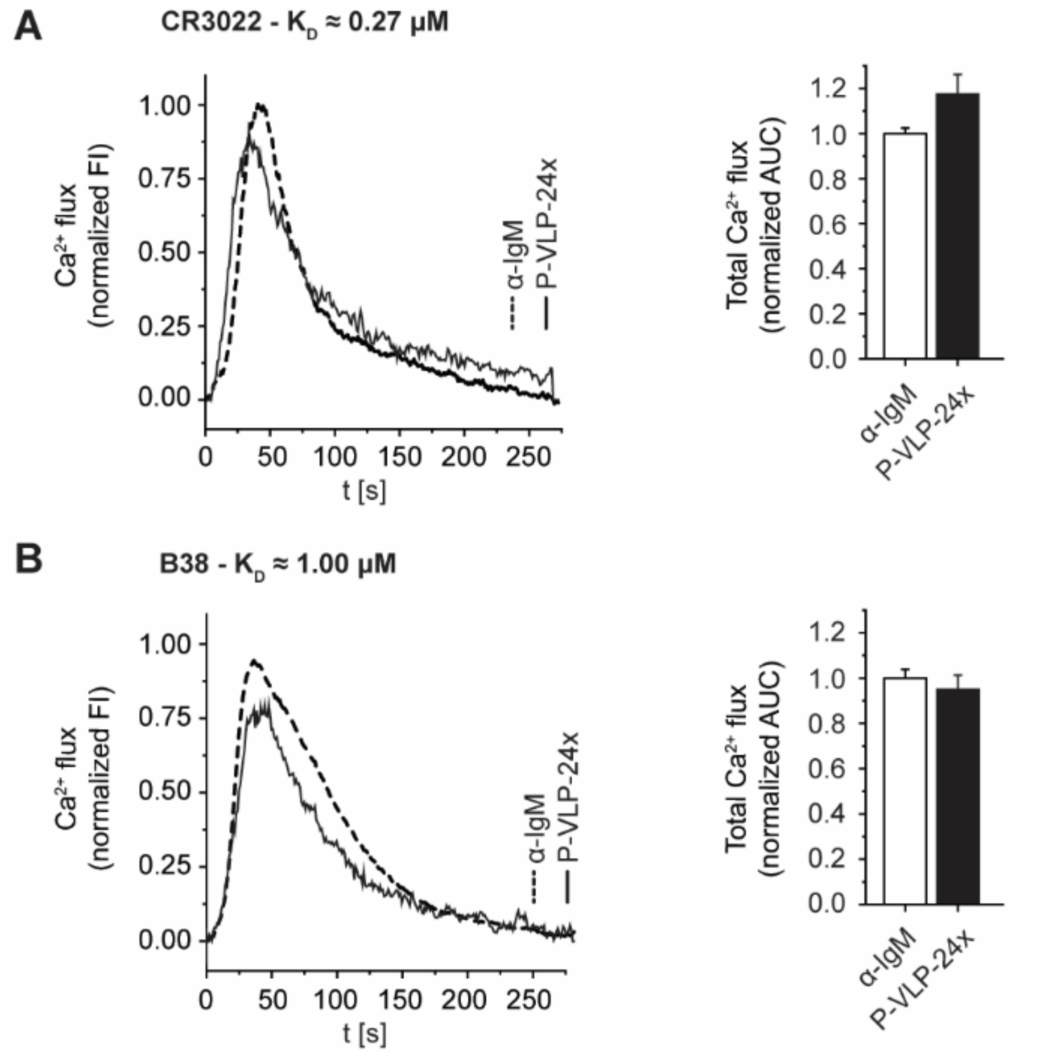


**Figure S7 – Supporting information for P-VLP B-cell activation.**
**(A and B)** Ramos B cells expressing the BCRs C3022 and B38 were incubated with α-IgM and **P-VLP-24x** at 30 nM RBD. $Ca^{2+}$ flux in response to RBD incubation was assayed using Fura Red. Representative fluorescence intensity curves are shown (left). Total $Ca^{2+}$ flux was quantified via the normalized AUC (right). Normalized AUC values were determined from n = 3 biological replicates. Error bars represent the standard error of the mean.

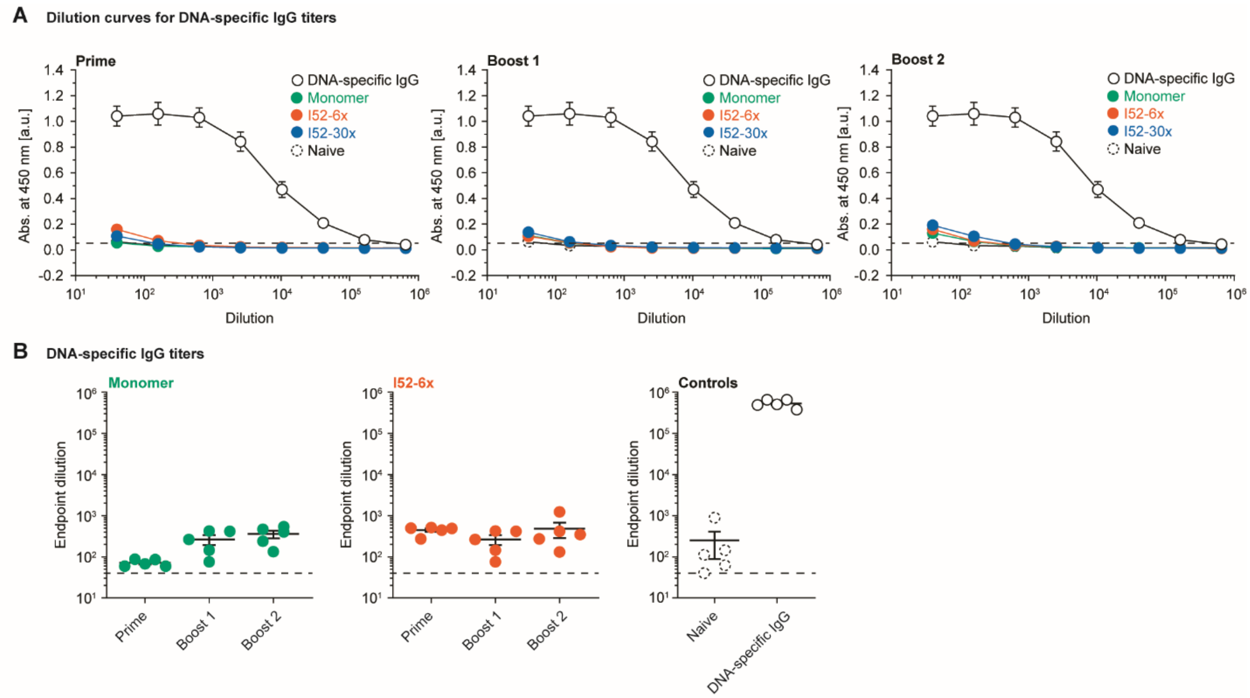


**Figure S8 – Supporting information for DNA-specific IgG ELISA**
**(A)** Dilution curves for DNA-specific IgG ELISAs with mouse sera following immunization with 7.5 µg RBD are shown. Plates were coated with calf thymus DNA. **(B)** DNA-specific IgG endpoint dilutions for monomer RBD and **I52-6x**. Naïve serum and DNA-specific IgG served as the positive and negative controls for the ELISA, respectively. DNA-specific IgG was diluted from 10 µg/ml. Absorbance values and IgG titers, were determined from n = 5 biological replicates. One-way ANOVA was performed followed by Dunnett’s T3 multiple comparison test at α = 0.05. Error bars represent the standard error of the mean.

### Supporting Tables

#### Table S1 – Scaffold sequence

AGAATTTCGCCATAGGTATGCTTAAGGAAGTCGAGATTGCGAACCATTACCGAGACTATGGCTTCATGTGGTGAT
TTCACCCGACCCACCCTTGGCGCCAGCTTTACGCAGCTTCCTGACGATACGTGGTGTAACGTTGTGTTTGGCAA
TGGAAACCGAGATCAACTATTTCTAATGCTGATATAGCAGAGTCTCGCGTCTATCATACGCAAGTCGCACGTCAT
TTTCGAGAGCAGCGTAAGACTCTGAAGGTCATGAGCCCAGATGTTATTACCCTCTACCTATAAACATCAAAATTG
TAGTCGTTTTACAGTCCATCGTCGCTCCAGAGCGAAGATTAAGGTTAGATCTAGATTATCTTTGCACGTGTGGAC
CGACGCAGCTGGGGCTCTAGCTCCACTACGGTTACGAAACTGCTGAACGATCTGGTCCACTTCAAGATTCACAC
ATCGTTTCATTCTTTGGACAACCAACACTCTCAGTCAGAGTTTCGAGTATAATAATTCTTCCGCGCTAGGGTAAAA
AGCAGATATGGGGAGACATTCCGGGCTTTTGAGCCGATACACTAAGCACTTGACATACTCACATCAGTAGAGGT
TAACATTCATGACTATCACGCGCTGCAGGAGCGCAACGCAATTAATGTGCGCCCTGTAGCGGCGCATTAAGCGC
GGCGGGTGTGGTGGTTACGCGCAGCGTGACCGCTACACTTGCCAGCGCCCTAGCGCCCGCTCCTTTCGCTTTC
TTCCCTTCCTTTCTCGCCACGTTCGCCGGCTTTCCCCGTCAAGCTCTAAATCGGGGGCTCCCTTTAGGGTTCCG
ATTTAGTGCTTTACGGCACCTCGACCCCAAAAAACTTGATTAGGGTGATGGTTCACGTAGTGGGCCATCGCCCT
GATAGACGGTTTTTCGCCCTTTGACGTTGGAGTCCACGTTCTTTAATAGTGGACTCTTGTTCCAAACTGGAACAA
CACTCAACCCTATCTCGGTCTATTCTTTTGATTTATAAGGGATTTTGCCGATTTCGGCCTATTGGTTAAAAAATGA
GCTGATTTAACAAAAATTTAACGCGAATTACAACCGGGGTACATATGATTGGGGTCTGACGCTCAGTGGAACGAA
AACTCACGTTAAGGGATTTTGGTCATGAGATTATCAAAAAGGATCTTCACCTAGATCCTTTTAAATTAAAAATGAA
GTTTTAAATCAATCTAAAGTATATATGAGTAAACTTGGTCTGACAGTTACCAATGCTTAATCAGTGAGGCACCTAT
CTCAGCGATCTGTCTATTTCGTTCATCCATAGTTGCCTGACTCCCCGTCGTGTAGATAACTACGATACGGGAGGG
CTTACCATCTGGCCCCAGTGCTGCAATGATACCGCGAGACCCACGCTCACCGGCTCCAGATTTATCAGCAATAA
ACCAGCCAGCCGGAAGGGCCGAGCGCATAAGTGGTCCTGCAACTTTATCCGCCTCCATCCAGTCTATTAATTGT
TGCCGGGAAGCTAGAGTAAGTAGTTCGCCAGTTAATAGTTTGCGCAACGTTGTTGCCATTGCTACAGGCATCGT
GGTGTCACGCTCGTCGTTTGGTATGGCTTCATTCAGCTCCGGTTCCCAACGATCAAGGCGAGTTACATGATCCC
CCATGTTGTGCAAAAAAGCGGTTAGCTCCTTCGGTCCTCCGATCGTTGTCAGAAGTAAGTTGGCCGCAGTGTTAT
CACTCATGGTTATGGCAGCACTGCATAATTCTCTTACTGTCATGCCATCCGTAAGATGCTTTTCTGTGACTGGTGA
GTACTCAACCAAGTCATTCTGAGAATAGTGTATGCGGCGACCGAGTTGCTCTTGCCCGGCGTCAATACGGGATA
ATACCGCGCCACATAGCAGAACTTTAAAAGTGCTCATCATTGGAAAACGTTCTTCGGGGCGAAAACTCTCAAGGA
TCTTACCGCTGTTGAGATCCAGTTCGATGTAACCCACTCGTGCACCCAACTGATCTTCAGCATCTTTTACTTTCAC
CAGCGTTTCTGGGTGAGCAAAAACAGGAAGGCAAAATGCCGCAAAAAAGGGAATAAGGGCGACACGGAAATGT
TGAATACTCATACTCTTCCTTTTTCAATATTATTGAAGCATTTATCAGGGTTATTGTCTCATGAGCGGATACATATT
TGAATGTATTTAGAAAAATAAACAAATAGGGGTTCCGCGCACATTTCCCCGAAAAGTGCCACCTGACGTCTAAGA
AACCATTATTATCATGACATTAACCTATAAAAATAGGCGTATCACGAGGCCCTTTCGTCGAATTCGTCGTCGTCCC
CTCAAACTCTTGGGTGGAGAGGCTATTCGTTTAAGGTCACATCGCATGTAATTTACTTATTCTCTGTTGTTGAGCC
ACCCGGGCGCCAGATTTTGTTTAAAGCTTTGTCTCTTAGTTTGTATAGACAGATTCAGAGTGCAAGGTTTCGTTC
GCTCGTACCTGGTTTTCCCTGGTTCTTCACAGATAGGATTTGACTTTCTACAACACTTATGCGGCTTCCTACCCGT
TTGAAGGCCGATACAGGTGCTGCGCAAAATGCGGGCGAACATAGAGTATCAAAACAACGCCTTCTAATCTAGGA
ATATAGGGAAGATACGTATTTGCTACCATGCTTTCTTGGGTCATTAACGACCAACCTCTTTTCTTTTAAAGTAGGA
TTGCACAATGAATGAATACACGTGGTCCGATAACTGACCAAGTAACATGGTTATCACTAGATGTCCGCCAGACGT
GTGCAAACCAACCCGGGAGTTACGTCACTAATCCTTCGCTACGTCGTGAAGATATTTACTTGTGAATATCGAGGG
TAATAAGATAATAGACTGTGACTAGTATTGCCAGACTGTCGCTACCTGCAACACATAACTATCCTGAGGTTACTG
CATAGTACTGATTACACCCGAGTCAAAATTTCTAACTTCTAACATGTACCTAGTAACCAGCTCAATAATTATGTCA
GAATATAGCTCTGGGAACCCTCGGACAATTATGATACACGGTATTAATATCTTGCTTGCGTTAGCCACTTCTCATC
TTTGGATACCGATTCTATTTTGCATAGCAGTTCCTTTTACACATATA

#### Table S2 – Staple sequences – I52

| Staple number | Sequence |
|---|---|
| 2 | ACATCTGGATGGCGAAATTCTTATATGTGGTAGAGGGTAATA |
| 3 | GCTCATGACCAAGCATACCT |
| 4 | GTTCGCAATCTTTTTTCGACTTCCTTTTCAGAGTCTTTTTTTACGCTGCTCT |
| 5 | CGGTAATGTCACCACATGAAGCCGTCCGAGG |
| 6 | GTTCCCAGACCGTGTATCATAATTATAGTCT |

| | |
|---|---|
| 7 | GCGAGACTCTGTTTTTCTATATCAGCCCAAACACAACTTTTTGTTACACCACCCAAGGGTG GGTTTTTTCGGGTGAAA |
| 8 | AGCTGGCGGTATCGTCAGGAAGCTAACCTCA |
| 9 | GGATAGTTAATCAGTACTATGCAGTGCGTAA |
| 10 | TTCCATTGATTAGAAATAGTTGAGTAGCGAA |
| 11 | GGATTAGTTAAATATCTTCACGACTCTCGGT |
| 12 | TGATAGACCGAAAATGACGTGCGACACGTGC |
| 13 | AAAGATAAAGCTGCGTCGGTCCACTTGCGTA |
| 14 | CTACAATTTTGTTTTTATGTTTATAGTAAAAGGAACTTTTTTGCTATGCAAATTTTTATAGGT TTTTTTAATGTCATG |
| 15 | GTCAAGTGCGACGATGGACTGTAAAACGACTGATGTGAGTAT |
| 16 | CTTAGTGTATCGCTCTGGAG |
| 17 | TCTAGATCTAATTTTTCCTTAATCTTCGGCTCAAAAGTTTTTCCCGGAATGT |
| 18 | AGAATTATTATTTTTTACTCGAAACTGAAACGATGTGTTTTTTGAATCTTGACCGTAGTGGA GTTTTTCTAGAGCCCC |
| 19 | TTTCGTAAAGTGGACCAGATCGTACATCTAG |
| 20 | TGATAACCTGCACACGTCTGGCGGTCAGCAG |
| 21 | CAAAGAATCTGACTGAGAGTGTTTGATAACA |
| 22 | CTGCGGCCCTGCCATAACCATGAGGGTTGTC |
| 23 | AGCGCGGACTCCCCATATCTGCTTACCCGCC |
| 24 | GCGCTTAAGCGCGTAACCACCACTTTACCCT |
| 25 | CAACGTCAAAGTTTTTGGCGAAAAACAGTCATGAATGTTTTTTTAACCTCTA |
| 26 | GATGGCCCTGCGCTCCTGCAGCGCGTGATCGTCTATCAGGGC |
| 27 | ACTACGTGAATTAATTGCGT |
| 28 | TGCGCCGCTACTTTTTAGGGCGCACACCATCACCCTATTTTTATCAAGTTTT |
| 29 | GAACCCTAAAGTTTTTGGAGCCCCCGCGTGGCGAGAATTTTTAGGAAGGGAAGGCAAGTG TAGTTTTTCGGTCACGCT |
| 30 | AGGGCGCTGAAAGCGAAAGGAGCTAACCGCT |
| 31 | TTTTTGCAGGAGGACCGAAGGAGCGGGCGCT |
| 32 | CCGGCGAAATTTAGAGCTTGACGATCATATG |
| 33 | TACCCCGGTGAGCGTCAGACCCCAGGGAAAG |
| 34 | CTAAATCGTTGGGGTCGAGGTGCCAGACCGA |
| 35 | GATAGGGTATAAATCAAAAGAATGTAAAGCA |
| 36 | CTTAGACGACTATTAAAGAACGTGGACTCATAATAATGGTTT |
| 37 | TCAGGTGGCAACAAGAGTCC |
| 38 | TGAGTGTTGTTTTTTTCCAGTTTGGACTTTTCGGGGATTTTTAATGTGCGCG |
| 39 | ATTCAAATATGTTTTTTATCCGCTCACGTAGTTATCTTTTTTACACGACGGGCCGAAATCGG CTTTTTAAAATCCCTT |
| 40 | ATGGATGAGCTCATTTTTTAACCAATAGGGAGTCAGGCAACT |
| 41 | ACGAAATAGATGTTAAATCA |
| 42 | TAACGTGAGTTTTTTTTTCGTTCCACTTGTAATTCGCTTTTTGTTAAATTTTCAGATCGCTGA TTTTTGATAGGTGCC |
| 43 | AAATCCCTCTTTTTGATAATCTCGAACCGGA |
| 44 | GCTGAATGTCGCCTTGATCGTTGGATGACCA |
| 45 | CAAGTTTACTCTTTTTATATATACTTAAAAGGATCTATTTTTGGTGAAGATC |
| 46 | TTTAATTTTAGATTGATTTAAAACTAGCTTC |

| | |
|---|---|
| 47 | CCGGCAACTGGCGAACTACTTACTCTTCATT |
| 48 | TGTCAGACTCACTGATTAAGCATGGAGCCGG |
| 49 | TGAGCGTGTATTGCTGATAAATCTTGGTAAC |
| 50 | CTGATAAAAGATGGTAAGCCCTCCCGTATTGAGACAATAACC |
| 51 | TGCTTCAATACACTGGGGCC |
| 52 | GGTCTCGCGGTTTTTTATCATTGCAGATATTGAAAAATTTTTGGAAGAGTATCCTTTTTTGC GTTTTTGCATTTTGCC |
| 53 | TGAAAGTAAAATTTTTGATGCTGAAGGGCCCTTCCGGTTTTTCTGGCTGGTT |
| 54 | ACGAGTGGTGCAGGACCACTTATGCGCTCATCAGTTGGGTGC |
| 55 | GTTACATCGACGGATAAAGT |
| 56 | AATTAATAGACTTTTTTGGATGGAGGACTGGATCTCATTTTTACAGCGGTAA |
| 57 | TTCCAATGATGTTTTTAGCACTTTTAATTGACGCCGGTTTTTGCAAGAGCAAAACGTTGCG CATTTTTAACTATTAAC |
| 58 | CAATGGCAACCTCGGTCGCC |
| 59 | GCCTGTAGGCATACACTATTCTCAGAATGGTGACACCACGAT |
| 60 | CAACATGGGGGTTTTTATCATGTAACAAGCCATACCATTTTTAACGACGAGCACTTGGTTG AGTTTTTTACTCACCAG |
| 61 | CAGTAAGAGAATTTTTTTATGCAGTGAACTTACTTCTTTTTTGACAACGATC |
| 62 | TGGCATGATCACAGAAAAGCATCTCATTGTG |
| 63 | CAATCCTAGACCACGTGTATTCATTTACGGA |
| 64 | TATCCCGTAAGTTCTGCTATGTGGGAAAGCA |
| 65 | TGGTAGCATCGTTAATGACCCAACGCGGTAT |
| 66 | AGAACGTTGATCCTTGAGAGTTTTCTTCAAA |
| 67 | CGGGTAGGAGCACCTGTATCGGCCGCCCCGA |
| 68 | AACGCTGGTTCCTGTTTTTGCTCGTGAAGAA |
| 69 | CCAGGGAAAAGTCAAATCCTATCTACCCAGA |
| 70 | CCTTATTCGAGTATTCAACATTTCTGGCTCA |
| 71 | ACAACAGAAATCTGGCGCCCGGGCGTGTCGC |
| 72 | CTAAATACGAACCCCTATTTGTTTAGCCTCT |
| 73 | CCACCCAATGTGACCTTAAACGAATATTTTT |
| 74 | CCAAAGATAAGGGCCTCGTGATACGCCTAATAGAATCGGTAT |
| 75 | GAGAAGTGGCAATTCGACGA |
| 76 | GAATAAGTAAATTTTTTTACATGCGAGAGTTTGAGGGTTTTTGACGACGACG |
| 77 | TAACGCAAGCATTTTTAGATATTAATAGCTATATTCTTTTTTGACATAATTAAAGAGACAAAG TTTTTCTTTAAACAA |
| 78 | TAGGTACACTGAATCTGTCTATACAAACTTTGAGCTGGTTAC |
| 79 | TGTTAGAAGTACCTTGCACT |
| 80 | AAGCCGCATAATTTTTGTGTTGTAGAAACCAGGTACGTTTTTAGCGAACGAA |
| 81 | TAGAAATTTTGTTTTTACTCGGGTGTATGTGTTGCAGTTTTTGTAGCGACAGATGTTCGCC CGTTTTTCATTTTGCGC |
| 82 | TTGATACTCTTCTGGCAATA |
| 83 | GCGTTGTTCTAGTCACAGTCTATTATCTTCCTAGATTAGAAG |
| 84 | ATGTTACTTGGTTTTTTCAGTTATCGCTTTAAAAGAATTTTTAAGAGGTTGGAATACGTATCT TTTTTTCCCTATATT |
| 85 | ATTACCCTCGATTTTTTATTCACAAGGACGTAACTCCTTTTTCGGGTTGGTT |

#### Table S3 – Modified staple sequences

**Cy5-modified staples**

| Staple number | Sequence |
| --- | --- |
| 19 | **/Cy5-TEG/TT**TTTCGTAAAGTGGACCAGATCGTACATCTAG |
| 30 | **/Cy5-TEG/TT**AGGGCGCTGAAAGCGAAAGGAGCTAACCGCT |
| 46 | **/Cy5-TEG/TT**TTTAATTTTAGATTGATTTAAAACTAGCTTC |
| 69 | **/Cy5-TEG/TT**CCAGGGAAAAGTCAAATCCTATCTACCCAGA |
| 72 | **/Cy5-TEG/TT**CTAAATACGAACCCCTATTTGTTTAGCCTCT |

**DBCO-modified staples – I52-1x-DBCO**

| Staple number | Sequence |
| --- | --- |
| 9 | **/DBCO-TEG/TT**GGATAGTTAATCAGTACTATGCAGTGCGTAA |

**DBCO-modified staples – I52-6x-DBCO**

| Staple number | Sequence |
| --- | --- |
| 9 | **/DBCO-TEG/TT**GGATAGTTAATCAGTACTATGCAGTGCGTAA |
| 15 | **/DBCO-TEG/TT**GTCAAGTGCGACGATGGACTGTAAAACGACTGATGTGAGTAT |
| 32 | **/DBCO-TEG/TT**CCGGCGAAATTTAGAGCTTGACGATCATATG |
| 54 | **/DBCO-TEG/TT**ACGAGTGGTGCAGGACCACTTATGCGCTCATCAGTTGGGTGC |
| 64 | **/DBCO-TEG/TT**TATCCCGTAAGTTCTGCTATGTGGGAAAGCA |
| 71 | **/DBCO-TEG/TT**ACAACAGAAATCTGGCGCCCGGGCGTGTCGC |

**DBCO-modified staples – I52-30x-DBCO**

| Staple number | Sequence |
| --- | --- |
| 2 | **/DBCO-TEG/TT**ACATCTGGATGGCGAAATTCTTATATGTGGTAGAGGGTAATA |
| 5 | **/DBCO-TEG/TT**CGGTAATGTCACCACATGAAGCCGTCCGAGG |
| 9 | **/DBCO-TEG/TT**GGATAGTTAATCAGTACTATGCAGTGCGTAA |
| 11 | **/DBCO-TEG/TT**GGATTAGTTAAATATCTTCACGACTCTCGGT |
| 13 | **/DBCO-TEG/TT**AAAGATAAAGCTGCGTCGGTCCACTTGCGTA |
| 15 | **/DBCO-TEG/TT**GTCAAGTGCGACGATGGACTGTAAAACGACTGATGTGAGTAT |
| 20 | **/DBCO-TEG/TT**TGATAACCTGCACACGTCTGGCGGTCAGCAG |
| 22 | **/DBCO-TEG/TT**CTGCGGCCCTGCCATAACCATGAGGGTTGTC |
| 24 | **/DBCO-TEG/TT**GCGCTTAAGCGCGTAACCACCACTTTACCCT |
| 26 | **/DBCO-TEG/TT**GATGGCCCTGCGCTCCTGCAGCGCGTGATCGTCTATCAGGGC |
| 31 | **/DBCO-TEG/TT**TTTTTGCAGGAGGACCGAAGGAGCGGGCGCT |
| 32 | **/DBCO-TEG/TT**CCGGCGAAATTTAGAGCTTGACGATCATATG |
| 35 | **/DBCO-TEG/TT**GATAGGGTATAAATCAAAAGAATGTAAAGCA |
| 36 | **/DBCO-TEG/TT**CTTAGACGACTATTAAAGAACGTGGACTCATAATAATGGTTT |
| 40 | **/DBCO-TEG/TT**ATGGATGAGCTCATTTTTTAACCAATAGGGAGTCAGGCAACT |
| 44 | **/DBCO-TEG/TT**GCTGAATGTCGCCTTGATCGTTGGATGACCA |
| 47 | **/DBCO-TEG/TT**CCGGCAACTGGCGAACTACTTACTCTTCATT |
| 48 | **/DBCO-TEG/TT**TGTCAGACTCACTGATTAAGCATGGAGCCGG |
| 50 | **/DBCO-TEG/TT**CTGATAAAAGATGGTAAGCCCTCCCGTATTGAGACAATAACC |
| 54 | **/DBCO-TEG/TT**ACGAGTGGTGCAGGACCACTTATGCGCTCATCAGTTGGGTGC |

| | |
|---|---|
| 59 | **/DBCO-TEG/TT**GCCTGTAGGCATACACTATTCTCAGAATGGTGACACCACGAT |
| 62 | **/DBCO-TEG/TT**TGGCATGATCACAGAAAAGCATCTCATTGTG |
| 64 | **/DBCO-TEG/TT**TATCCCGTAAGTTCTGCTATGTGGGAAAGCA |
| 67 | **/DBCO-TEG/TT**CGGGTAGGAGCACCTGTATCGGCCGCCCCGA |
| 68 | **/DBCO-TEG/TT**AACGCTGGTTCCTGTTTTTGCTCGTGAAGAA |
| 71 | **/DBCO-TEG/TT**ACAACAGAAATCTGGCGCCCGGGCGTGTCGC |
| 73 | **/DBCO-TEG/TT**CCACCCAATGTGACCTTAAACGAATATTTTT |
| 74 | **/DBCO-TEG/TT**CCAAAGATAAGGGCCTCGTGATACGCCTAATAGAATCGGTAT |
| 78 | **/DBCO-TEG/TT**TAGGTACACTGAATCTGTCTATACAAACTTTGAGCTGGTTAC |
| 83 | **/DBCO-TEG/TT**GCGTTGTTCTAGTCACAGTCTATTATCTTCCTAGATTAGAAG |
